# The pleuroparenchymal fibroelastosis atlas reveals aberrant cell states and their zonation as an alternate roadmap to lung fibrosis

**DOI:** 10.1101/2025.07.06.663336

**Authors:** Jannik Ruwisch, Aurélie Cazes, Lena M Leiber, Raphael Borie, Lavinia Neubert, Christian Leonard, Vincent Thomas de Montpréville, Adam Szmul, Farah Moussa, Stijn E Verleden, Svenja Gaedcke, Jan Hegermann, Jan Fuge, Matthias Ballmaier, Jan C Kamp, Mark Greer, Peter Braubach, Christopher Werlein, Fabius Ius, Theresa Graalmann, Kalil Aburahma, Laurens J De Sadeleer, Ryoko Egashira, Maximilian Ackermann, Daisuke Yamada, Marius M Hoeper, Christine Falk, Jens Gottlieb, Herbert B Schiller, Bart Vanaudenaerde, Benjamin Seeliger, Marie Pierre Debray, Jean Francois Bernaudin, Lars Knudsen, Emmanuel Bergot, Joseph Jacob, Hervé Mal, Danny Jonigk, Sabine Dettmer, Pierre Mordant, Antje Prasse, Elie Fadel, Wim A Wuyts, Bruno Crestani, Naftali Kaminski, Aurélien Justet, Jonas C Schupp

## Abstract

**Background:** Pleuroparenchymal fibroelastosis (PPFE) is a progressive interstitial lung disease with higher prevalence in females, histologically characterized by intra-alveolar fibrosis with septal elastosis (AFE). Effective treatments are lacking, highlighting the need to dissect its pathogenesis at single-cell resolution.

**Methods:** We performed single-nucleus RNA sequencing (snRNAseq) on explanted lungs from a German (n=23) and a French cohort (n=17) of PPFE patients, and controls (n=16). Identified cell populations were localized by immunofluorescence and multiplex RNA *in-situ* hybridization. Hierarchical phase-contrast computed tomography (HiP-CT) and micro-CT provided 3D spatial context. Reanalyzed snRNAseq data from a Belgian IPF cohort (n=9) served as disease comparator.

**Findings:** We present the first snRNAseq atlas of PPFE’s cellular and structural landscape based on a European multinational cohort. 24 PPFE patients were female (60.0%), while 34 were non-smokers (85.0%). 519,920 nuclear transcriptomes from PPFE and IPF patients, and controls were profiled. We identified PPFE-specific accumulations of *MFAP5*+*PI16*+*SFRP2*+ adventitial and *LEPR*+*ITGA8*+*SFRP2*+*DIO2*+ elastofibrotic fibroblasts as main drivers of elastotic remodeling in PPFE. Multiple PPFE fibroblast subsets acquire an inflammatory activation state as highlighted by the expression of *CXCL12* and *CXCL14.* This is accompanied by a marked increase in lymphocytes and the formation of tertiary lymphoid structures (TLS) in a disease that was previously considered to be purely elastofibrotic. We identified *CTHRC1*+ fibrotic fibroblasts and Aberrant Basaloid cells in PPFE as well, forming the “Usual Fibrotic Niche”. 3D reconstruction of the pronounced *COL15A1*+ vascular conglomerate at the border of the elastofibrotic and subpleural fibrosis indicates communication with interlobar veins. Last, we observed a zonation of the PPFE lesion, constructed by the above-mentioned PPFE-associated cell types.

**Interpretation:** Our unprecedented cellular and molecular survey uncovers previously unobserved PPFE-specific inflammatory and elastogenic fibroblast populations, as well as the presence of *CTHRC1*+ fibroblasts and Aberrant Basaloid cells common to other fibrotic ILDs. These findings provide the foundation for including PPFE patients in current antifibrotic trials, as well as development of PPFE-specific therapies.

**Funding:** Supported mainly by the Else Kröner-Fresenius Foundation, the German Center for Lung Research and the Fondation du Souffle.

## Introduction

Pleuroparenchymal fibroelastosis (PPFE) is a rapidly progressive interstitial lung disease (ILD) that predominantly affects females^1–3^. Aside from lung transplantation, no specific treatment options are available for PPFE^4^. In most cases, the etiology of PPFE remains idiopathic (iPPFE). PPFE can occur secondary to allogenic stem cell transplantation, autoimmune conditions or lung transplantation, in the latter case referred to as restrictive allograft syndrome (RAS). PPFE histopathology is characterized by dense fibrosis of the visceral pleura and the pattern of intra-alveolar fibrosis with septal elastosis (AFE) with deposition of elastic fibers along the residual alveolar walls and intra-alveolar deposition of collagen fibrils^1,5,6^. In addition, lymphoid cell aggregates have been frequently observed in AFE lesions^5^.

The advent of single cell RNA sequencing (scRNAseq) has dramatically revised our understanding of biology in general^7–9^ and respiratory disease pathogenesis in particular.^10–14^ This is especially true for idiopathic pulmonary fibrosis (IPF)^10,13^, the most common ILD^15,16^. IPF is characterized by a unique histological pattern called Usual Interstitial Pneumonia (UIP)^17^. scRNAseq enabled the identification of aberrant cell populations that build the UIP pattern of IPF including *CTHRC1*+ fibrotic fibroblasts, Aberrant Basaloid cells, ectopic systemic-venous endothelial cells and *SPP1*+ profibrotic macrophages^10^. However, scRNAseq efforts in fibrotic ILDs have focused predominantly on UIP, leaving a knowledge gap for non-UIP ILDs.

In addition, the transferability of scRNAseq findings to other fibrotic ILDs such as PPFE remains uncertain as IPF and PPFE are radiologically and histologically reflected by different morphological processes^18^: PPFE typically affects non-smoking female patients with relatively young age^1,2,4^, while IPF occurs more commonly in aged male ex-smokers^1,4,16^. PPFE usually affects the upper lobes of the lungs, while IPF remodeling predominates in the lower lobes. Histopathologically, UIP is characterized by spatiotemporal heterogenous fibrosis, destruction of the lung architecture and bronchiolization of the distal lung^17^, while AFE is a homogenous fibrosis with preserved lung scaffolds and global lung architecture^5,6^.

In view of these substantial differences between PPFE and IPF, the following major knowledge gaps exist: it is not known whether PPFE is also composed of the usual IPF-associated pathological cell types, or whether PPFE-specific cell populations exist. And more importantly, which common disease-spanning principles of lung fibrosis can we derive by studying two prototypically progressive but clinically divergent ILDs? Lastly, PPFE research has lacked a compass to guide which cellular or molecular targets would be useful to test in clinical trials. Closing these knowledge gaps is essential to develop ILD-spanning treatment approaches and for providing a rationale for PPFE-specific treatment approaches.

Here, we set up the first multinational single-cell-resolved PPFE atlas. Following integration with recently published IPF dataset (GSE286182)^19^, we capture the cellular landscape of these progressively fibrotic ILDs. For validation and localization purposes, we employ micro-CT guided multiplex RNA *in-situ* hybridization (ISH), immune fluorescence microscopy (IF), hierarchical phase-contrast (HiP) synchrotron CT^20^ and transmission electron microscopy (TEM), and complement these with extensive *in-silico* ligand-receptor analysis.

## Results

### Cohort and dataset characteristics

Biobanked lung tissue from 40 PPFE patients undergoing lung transplantation in three European ILD and transplant centers (Hannover, Germany, PPFE-GER; n=23, and two in Paris, France, PPFE-FR; n=17) as well as 16 controls (**Fig. 1A**, **table E1**) were analyzed. PPFE diagnosis was confirmed based on HRCT and histopathology in multidisciplinary team discussion (**table E3, Fig. S1, Fig. S2**). Female patients were more prevalent in both PPFE cohorts^21,22^ (PPFE-GER: 65.2%; PPFE-FR: 52.9%, **table E3**). The median age at diagnosis was 41.3 years [Q1=29.3, Q3=50.8 years], while prevalence of tobacco smoking was limited to 15% in the PPFE cohort. Our PPFE cohorts are therefore dominated by young, non-smoking and female patients. This contrasts the demographics of published IPF scRNAseq datasets which consisted mainly of male ex-smokers with advanced age^10,13^. The etiology of PPFE had remained idiopathic in most cases, despite extensive diagnostic work-up (PPFE-GER 87.0%; PPFE-FR 82,4%, **table E3**).

**Figure 1.**
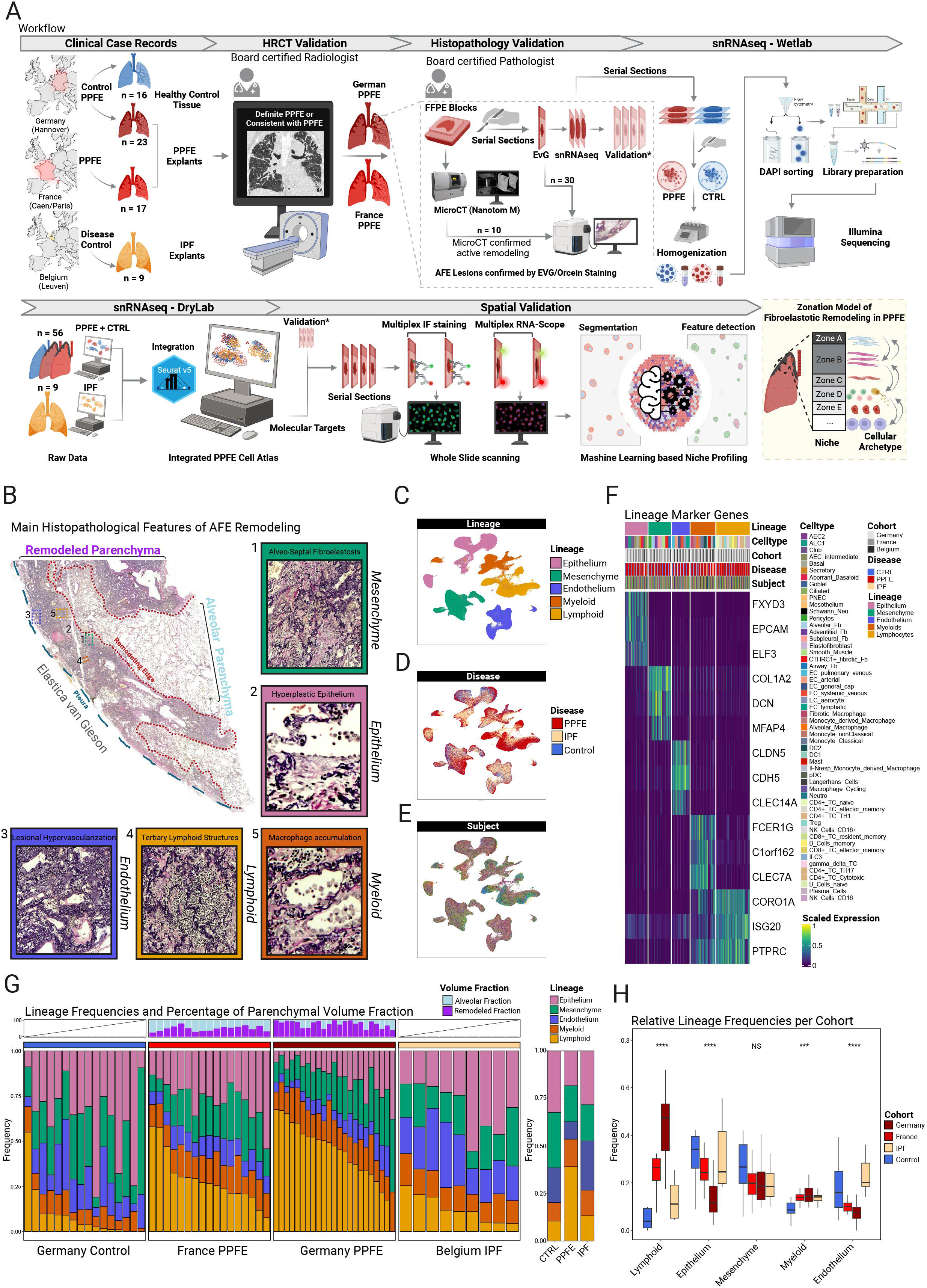
Experimental design and analysis of lineage composition. **A** Formalin-fixed and paraffin-embedded lung tissue from 23 German and 17 French patients was utilized to construct the 320,541 single nuclei spanning PPFE atlas, while published snRNAseq data of 199,379 nuclei from 9 IPF patients (GSE286182) served as disease control. Radiologic PPFE pattern and histological AFE pattern was confirmed by board-certified radiologist and pathologist, respectively. Micro-CT scanning was performed on FFPE blocks from 10 patients for targeted sampling of sites with postulated ongoing remodeling. 1 FFPE block per PPFE and per CTRL lung was serially sectioned. Initial 50µm thick sections were used for nuclei isolation, while subsequent 5µm sections served for validation via immune fluorescence staining or RNA in-situ hybridization. Automated whole slide cell segmentation in conjunction with artificial neural network (ANN) algorithm-based cell classification was used to for spatial niche profiling. **B** *Elastica van Gieson* stained PPFE slides can be segmented into remodeled (purple) and un-remodeled alveolar parenchyma (light blue). Both zones are demarcated by a sharp line referred to as (fibro-elastotic) remodeling edge. Within the remodeled parenchyma PPFE lesions typically consist of subpleural eosinophilic fibrosis and adjacent areas of alveolar fibroelastosis (AFE). Colored boxes highlight lineage related histopathological hallmarks of AFE remodeling. **C** Uniform Manifold Approximation and Projection (UMAP) representation of 519,920 nuclei, color-coded according to lineage. **D** UMAP colored by disease, with pooled color coding for the German and French PPFE cohorts. **E** UMAP with unique color coding for each investigated subject. **F** Unity scaled average expression of canonical lineage markers per celltype per subject. Subsamples were grouped after linage, celltype, cohort and disease. Each subject was assigned to a unique color. **G** Stacked bar plots of relative lineage frequencies per subject colored by lineage and split by cohort. Subjects were ordered after frequencies of lymphoid cells (lower panel). Surface fractions were calculated for remodeled (light blue) and un-remodeled (purple) tissue segmented on serially sectioned EvG stained images as exemplified in B (top panel). **H** Relative cell frequencies have been calculated as average per subject and are presented as a stacked bar plot. Illustrations were created with biorender.com.

In total, 320,541 single nuclei transcriptomes were profiled. We integrated our dataset with 199,379 nuclei sampled from explant tissue of nine IPF patients from Leuven, Belgium (**Fig. 1A**, GSE286182^19^). All major cell types of the human lung were identified based on distinct marker gene expression (**Fig. 1C, 1F**). Cell frequencies by lineage and by cell type per disease are provided in the supplemental results, and selected conspicuities are highlighted below. Raw sequencing data of the PPFE atlas will be deposited on Gene Expression Omnibus, processed data can be explored on the PPFE atlas webpage (PPFEatlas.com).

### Mesenchymal diversity in PPFE

Mesenchymal cells are main contributors to collagen and elastic fiber deposition in PPFE and constitute 22.8% of all profiled cells. *CTHRC1*+ fibrotic fibroblasts - the major profibrotic cell type in the IPF lung^10,21^ with high expression of collagen synthesis related genes (*COL1A1*, *COL3A1*) - occurred not only in IPF (9.18±4.11%, adj.pval_vs.CTRL_<0.001) but to a similar extent in the PPFE lung (PPFE-GER: 9.18±3.46%, adj.pval_vs.CTRL_<0.001 and PPFE-FR: 9.19±3.83% adj.pval_vs.CTRL_<0.001, **table E11**). ssGSEA for matrisomeDB^22^-related extracellular matrix (ECM) domains reassured *CTHRC1*+ fibrotic fibroblasts as the main source of parenchymal collagen and proteoglycans such as *DCN*, *BGN* and *LUM* in the PPFE lung (**Fig. 2L**).

**Figure 2.**
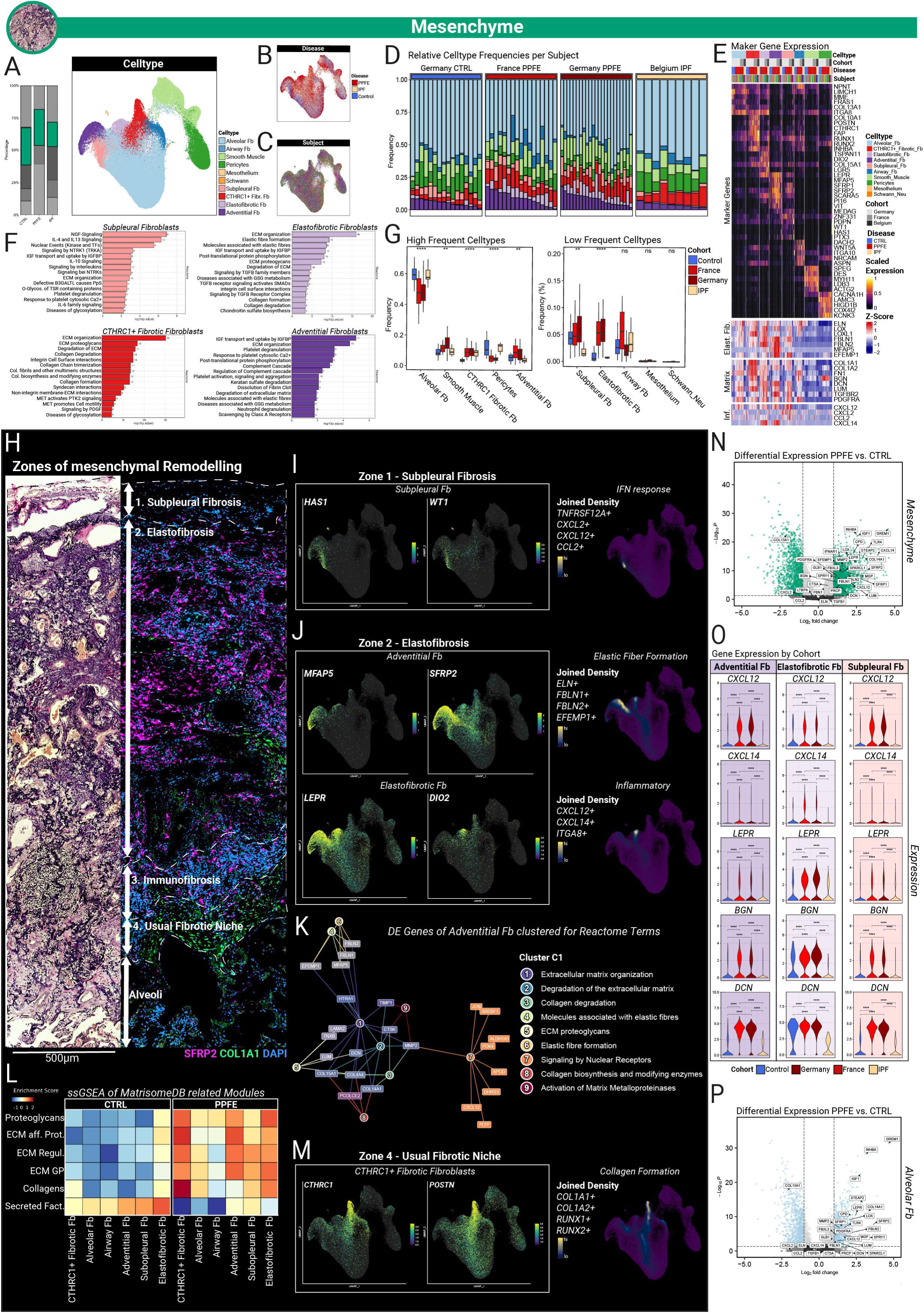
Mesenchymal archetypes in the PPFE lung. **A** Relative fraction of mesenchymal nuclei and UMAP embedding of identified mesenchymal cell types (Alveolar fibroblasts, airway fibroblasts, smooth muscle cells, pericytes, mesothelial cells, Schwann cells, subpleural fibroblasts, CTHRC1+ fibrotic fibroblasts, elastofibrotic fibroblasts and adventitial fibroblast) within the mesenchymal lineage **B** UMAP embedding by disease. **C** UMAP embedding by subject **D** Stacked bar plots of celltype frequencies per subject colored by celltype and split by cohort. Subjects were ordered after frequencies of adventitial fibroblasts. **E** Unity scaled average expression of canonical markers per subject (upper heatmap). Subjects were grouped after cohort and disease. Each subject was assigned to a unique color. Genes related to elastic fiber formation, extra cellular matrix remodeling and inflammatory signaling were z-score normalized per average expression per celltype and subject and plotted across cohorts and disease (lower heatmap). **F** Reactome Pathways based enrichment analysis of significantly differentially expressed genes of distinct fibroblasts populations. Number behind the bar indicates number of enriched genes per Reactome term. **G** Boxplots display the relative distribution of each mesenchymal cell type relative to the total mesenchymal lineage compartment, stratified by disease-related cohort. Whiskers indicate 1.5 times the interquartile range (IQR). Statistical comparisons between PPFE-GER, PPFE-FR, CTRL and IPF were performed using Kruskal-Wallis test with *: p ≤ 0.05, **: p ≤ 0.01, ***: p ≤ 0.001, and ****: p ≤ 0.0001. Post-hoc tests were performed under use of the Wilcoxon Rank Sum test, with Bonferroni correction for multiple comparisons across cohorts and celltypes. Adjusted p-values less than 0.05 were considered statistically significant. Detailed statistics are provided in **table E11**. **H** Correlative *Elastica van Gieson* and RNA *in-situ* hybridization-based stains of different pathogenic fibroblast archetypes in PPFE. Serial sections illustrate spatial zonation of remodeling in PPFE. **I** Featureplots of signature genes expressed by subpleural fibroblasts. **J** Featureplots of selected genes expressed in either adventitial or elastofibrotic fibroblasts. **K** Clustermap of enriched Reactome terms. Reactome terms were clustered with the greedy cluster algorithm by shared gene signature. Nodes represent count of enriched genes per term, while edges represent the intersection of enriched genes among terms (left panel). Genes enriched for terms (1-9) in cluster C1 are highlighted in the feature map (right panel). Clusters C2-C5 are provided in **Fig. S4** and **table E27**. **L** single sample gene set enrichment analysis (ssGSEA) for MatrisomeDB curated ECM gene modules in mesenchymal celltypes. Enrichment scores were calculated on cell level and subsequently aggregated per celltype. Heatmap values were z-score normalized to highlight differences across celltypes and diseases. **M** Feature plots of selected genes expressed in CTHRC1+ fibrotic fibroblasts. **N** Volcanoplot of differentially expressed genes following DESeq2 based pseudobulk aggregation between PPFE and CTRL in the mesenchymal lineage. P values were Bonferroni corrected for multiple comparisons across all tested genes. Genes exceeding log2 fold change (log2FC) > 1 and adj. pval < 0.05 were colored in green. Supervised gene annotation highlights genes related to remodeling in PPFE. **O** Differential gene expression of *CXCL12*, *CXCL14*, *LEPR* and *BGN* between cohorts in adventitial, elastofibrotic and subpleural fibroblasts. Plot background color corresponds the respective celltype related color code used in A. Violinplot color indicates cohort color. Inter-cohort statistics were assessed on cell level under use of Wilcoxon-Rank sum test with *: p ≤ 0.05, **: p ≤ 0.01, ***: p ≤ 0.001, and ****: p ≤ 0.0001. **P** Volcanoplot of differentially expressed genes following DESeq2 based pseudobulk aggregation between PPFE and CTRL in alveolar fibroblasts. P values were Bonferroni corrected for multiple comparisons across all tested genes. Genes exceeding log2FC>1 and adj. pval<0.05 were colored in green. Gene annotation was performed in a supervised fashion, highlighting genes related to remodeling in PPFE.

The two most PPFE-specific intra-mesenchymal shifts, however, were noted for adventitial fibroblasts (PPFE-GER: 9.04±5.15%, adj.pval_vs.CTRL_=0.023 and PPFE-FR 8.19%±4.13%, adj.pval_vs.CTRL_=0.024) and adventitial-like “elastofibrotic” fibroblasts^23^ (**Fig. 2G, table E11**). The latter received their name due to their transcriptional overlap with adventitial (*SFRP2*, *SCARA5*, *PI16*) and *CTHRC1*+ fibrotic fibroblasts (*INHBA*, *FN1, ASPN,* **Fig. 2E**). In addition, elastofibrotic fibroblasts co-expressed *LEPR* and the alveolar fibroblast marker *ITGA8*^24,25^, while lacking the adventitial fibroblasts marker *MFAP5 (***Fig. 2E,J**). Particularly, *LEPR* expression emphasizes the elastotic-to-profibrotic fate plasticity of elastofibrotic fibroblasts, as murine *Lepr*+ fibroblasts represent the major precursor population of *Cthcr1*+ fibrotic fibroblasts in mice^24^. Additional discriminating marker genes of elastofibrotic fibroblasts included *DIO2*, *COL15A1* and lipid metabolism related genes such as *APOD* or *STEAP2*, **table E12**, **Fig. 2J**).

Further characterization of PPFE enriched adventitial fibroblasts by differentially expressed gene enrichment analysis identified four clusters of pathways (**Fig. 2K, Fig. S3A**). Cluster C1 comprised ECM related genes involved in elastic fiber metabolism (*EFEMP1*, *FBLN2*, *FBLN1*, *MFAP5*) and collagens (*COL4A4*, *COL15A1*, *COL14A1,* **Fig. 2K**), while cluster C3 depicted an induction of glycosaminoglycan metabolism (**Fig. S4A**). Accordingly, we observed increased expression of charged glycosaminoglycans like small leucine-rich proteoglycans *BGN* and *DCN* in PPFE that are required for extracellular fibril formation of elastic fibers^26,27^ (**Fig. 2O**). Cluster C4 showed induced expression of genes related to dissolution of fibrin clots (*COLEC12*, *FTL, SCARA5*) whose deposition has been considered to precede the formation of AFE lesions^5^, while cluster C2 was involved in pro-metabolic signaling of insulin-like growth factors (*IGF1*, *IGFB6*, *VEGFB*, *CSF1*, **Fig. 2K**). This latter enrichment was complemented with transcriptional upregulation of the elastic fiber upstream regulator *IGF1* in PPFE mesenchyme compared to CTRL (**Fig. 2F**).

Notably, PPFE alveolar fibroblasts exhibited a significantly increased expression of *GREM1* and *IGF1* as well as of the adventitial collagen *COL14A1*, which may illustrate an early phenotypic shift from resident alveolar cell populations to mesenchymal cellular archetypes physiologically located in the loose connective tissue of the interlobar septum, broncho-vascular bundle or the subpleural connective tissue (**Fig. 2P**)^11^.

### Inflammatory activation states of fibroblasts in PPFE

Intra-mesenchymal differential expression analysis between PPFE and CTRL uncovered upregulation of a gene battery characteristic for a recently described mesenchymal cell state termed “inflammatory fibroblasts”^28^ including *CXCL12* and *CXCL14* as well as interferon response genes such as *IFNAR1* (**Fig. 2N**). In murine bleomycin-induced lung fibrosis, *Cxcl12+* inflammatory fibroblasts arise from alveolar fibroblasts and differentiate into *CTRHC1*+ fibrotic fibroblasts^25^, while localizing to sites of fibrotic remodeling in IPF^25^. In PPFE, multiple fibroblast subsets including subpleural, adventitial and elastofibrotic fibroblasts acquire this inflammatory activation state as highlighted by the expression of *CXCL12*, a known chemoattractant of lymphocytes (**Fig. 2O**). However, the details of the inflammatory activation vary: *HAS1*+*WT1*+ subpleural fibroblasts exclusively expressed an interferon/TNF-response signature with co-expression of *CXCL12*, *CXCL2*, *CCL2* and *TNFRSF12A (***Fig. 2I**). Co-expression of *CXCL12* and *CXCL14* was restricted to *LEPR*+*ITGA8*+*DIO2*+ elastofibrotic fibroblasts and *SFRP2*+ adventitial fibroblasts (**Fig. 2J**). In contrast to PPFE, *CXCL14* and *CXCL12* expression in mesenchymal cells from IPF or CTRL samples was almost absent (**Fig. 2O**).

### PPFE remodeling is spatially organized into four distinct zones

Spatial *in-situ* mapping of mesenchymal marker genes revealed a spatially organized disease pattern in PPFE, defining four niches with distinct cellular composition (**Fig. 2H**). From the pleural surface to apparently unremodeled alveolar parenchyma, these zones comprised: (**A**) a “Subpleural Fibrosis Zone”, (**B**) a zone marked by septal elastosis and intra-alveolar collagen deposition referred to as “Elastofibrosis Zone”, (**C**) an “Immunofibrotic Zone” and (**D**) a continuous band of *CTRHC1*+ fibrotic fibroblasts overlined by a monolayer of Aberrant Basaloid cells at the fibro-alveolar remodeling edge, a composition known from fibroblast foci in IPF. We therefore coined their joint emergence despite a different histological gestalt “Usual Fibrotic Niche”.

In the subpleural fibrosis zone, *HAS1*+ subpleural fibroblasts^13^ were identified and localized to a narrow area adjacent to the likewise *HAS1*+ mesothelium (**Fig. 3A-C**). On hematoxylin-eosin and EvG stains, this area corresponded to highly eosinophilic regions with low cellularity and abundant ECM deposition (**Fig. 1B**, **Fig. 2H**). The extent of this zone varied highly among patients in both cohorts and seemed to only serve as poor indicator of overall affected tissue within the specimen (**Fig. 3M**, blue bar, **S3**). Notably, we occasionally observed aggregates of CD4+, CD8+ and CD20+ lymphocytes in the Subpleural Fibrosis Zone (**Fig. 3N**). These aggregates often maintained a highly organized structure and are referred to as tertiary lymphoid structures (TLS: non-capsulated organized lymphoid cell aggregates in non-lymphoid tissues^29,30^). Notably, *CXCL12* chemotaxis is sufficient to induce TLS formation^31^, and *CXCL12*+ inflammatory cancer-associated fibroblasts co-localized with TLS in the fibrotic tumor environment^32^. Indeed, *CXCL12* expression was spatially enriched in the Subpleural Fibrosis Zone in proximity to TLS (**Fig. 3D**). On single cell level, this is paralleled by the high expression of *CXCL12* expression in *HAS1*+ subpleural fibroblasts (**Fig. 2O**).

**Figure 3.**
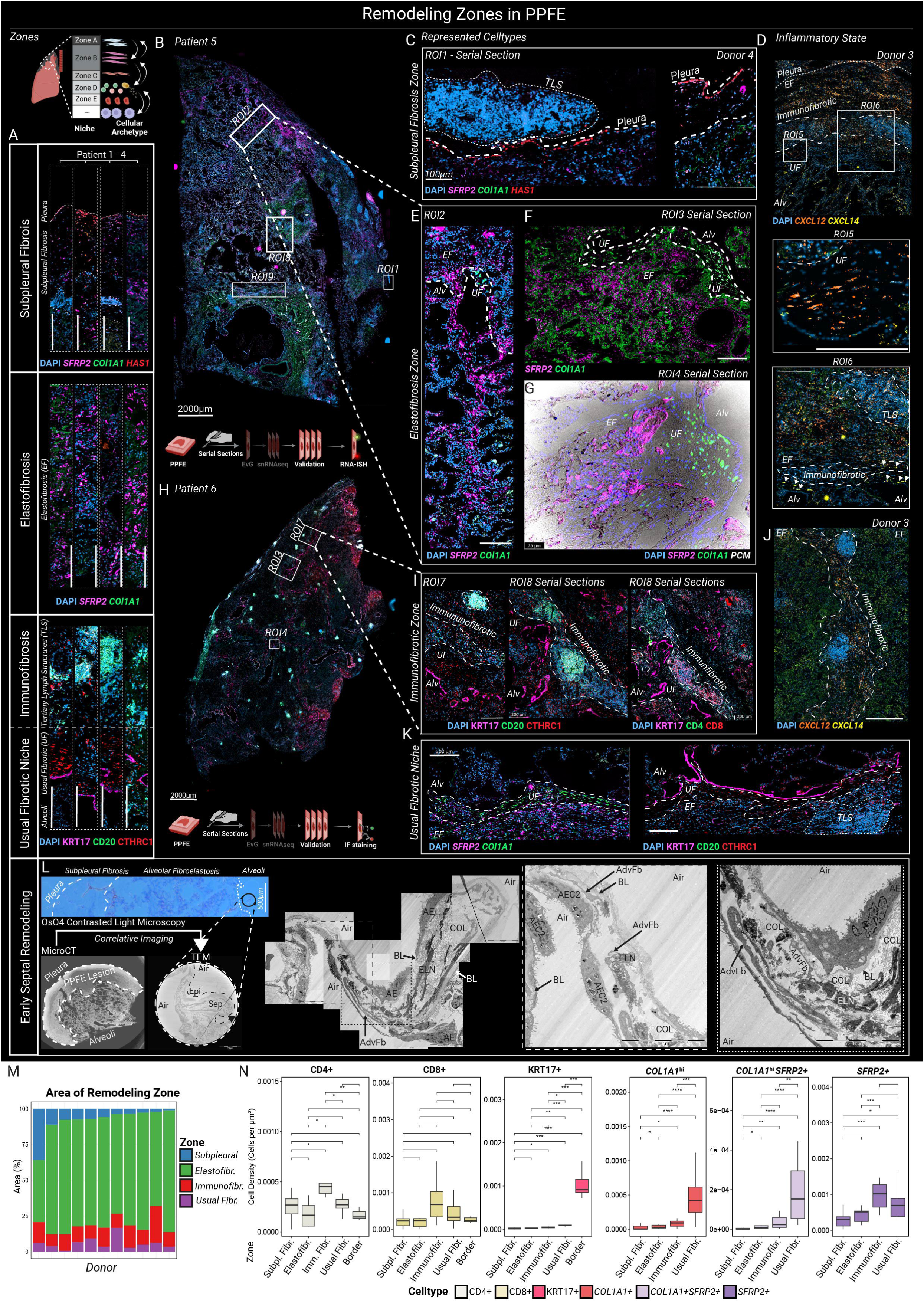
Zonation in the PPFE lung. **A** Overview of the major zones of remodeling in PPFE indicated by IF and ISH. Each box array of images represents a distinct donor. The pleural surface is indicated by a dashed line. **B** ISH of upper lobe PPFE lesion at low magnification. Boxes mark regions of interest (ROI) that are shown in higher magnification. **C** Subpleural Fibrosis Zone. High magnification of ISH staining of the Subpleural Fibrosis Zone. **D** Spatial expression of inflammatory fibroblast-related chemokines. Boxes indicate higher magnification of the Usual Fibrotic niche (ROI5) and the remodeling edge (ROI6). Pleura (fine) and Immunofibrotic Zone (coarse) are highlighted by a dashed line. Arrows indicate *CXCL14*+ cells within the Usual Fibrotic Niche **E** Elastofibrosis Zone. High magnification of septal remodeling within the aerated alveolar parenchyma. The remodeling edge is highlighted by a dashed line. **F** High magnification of ISH stained lesions of elastofibrosis adjacent to the Usual Fibrotic Niche (dashed line). **G** High magnification image of the transition zone between elastofibrosis and Usual Fibrotic Niche. Phase contrast (PCM) overlay highlights elastic fibers. **H** IF of upper lobe PPFE lesion at low magnification. Boxes mark regions of interest (ROI) that are shown in higher magnification. **I** Immunofibrotic Zone. High magnification IF images of tertiary lymphoid structure forming from diverse B and T cell subsets. **J** Spatial expression of inflammatory fibroblast chemokines in the immune cell niche. **K** Usual Fibrotic niche(dashed line). IF and ISH staining on serial sections of the Usual Fibrotic niche. Immune cell aggregates are encircled by a dashed line. **L** Micro-CT guided correlative imaging of early septal thickening beyond the fibro-alveolar remodeling edge. Scale bars resemble 20µm (low mag) and 10µm (high mag). Dashed boxes indicated areas of interest for higher magnification imaging. Fibroblast with adventitial fibroblast like morphology (Adv. Fb), alveolar epithelium (AE), basal lamina (BL), alveolar epithelial cells type 2 (AEC2), collagen (COL) and elastin (ELN) are annotated accordingly. **M** Stacked bar plots of manually segmented remodeling zones in 11 investigated PPFE-GER specimens referenced to the total of remodeled parenchyma. **N** Calculated cell densities of artificial neural network classified celltype based on multiplex ISH and IF staining in each remodeling zone. Statistical comparison among zones was performed under use of Wilcoxon rank sum test with *: p ≤ 0.05, **: p ≤ 0.01, ***: p ≤ 0.001, and ****: p ≤ 0.0001. Scale bars in ISH and IF stains represent 200µm unless stated otherwise. Illustrations were created with biorender.com

Below, in the Elastofibrosis Zone, we located *SFRP2+* adventitial and elastofibrotic fibroblasts along remnants of remodeled alveolar septa, and within them (**Fig. 3E,G**). RNA ISH coupled with phase-contrast microscopy located *SFRP2+* fibroblasts in close proximity to deposited elastic fibers within established AFE lesions and in thickened alveolar septa next to AFE lesions (**Fig. 3A,E-G)**. Micro-CT guided ultrastructural analysis of such thickened alveolar septa illustrated expanded interstitial collagen deposition and elastic fiber formation (**Fig. 3L**). Surprisingly, expansion of the interstitium correlated with a continuous phenotypic alteration of the overlining alveolar epithelium with loss of lamellar bodies and increasing thickening of the interposed basal lamina on correlative ultrastructural examination (**Fig. 3L**).

The Immunofibrotic Zone, an organized band of immune cell infiltrates, separates the Elastofibrosis Zone from the Usual Fibrotic Niche (**Fig. 3A**,**I**). The relative extent of this immune cell layer was well conserved among all histologically investigated PPFE specimen (n=11, **Fig. 3M**). Semi-supervised cell quantification revealed highest densities for CD4+ and CD8+lymphocytes in this zone in concordance with frequently observed TLS composed of CD4+, CD8+ and CD20+ lymphocytes (**Fig. 3A,I**). Furthermore, afferent and efferent vasculature from which immune cells are recruited are involved in TLS formation^29,30^. Accordingly, we frequently observed *COL15A1*+ *PLVAP*+ as well as *PDPN*+ lymphatic vasculature associated with formed TLS. Remarkably, inflammatory *SFRP2*+ fibroblasts were most abundant in the Immunofibrotic Zone as well (**Fig. 3N**), highlighting the local immuno-mesenchymal crosstalk. In fact, we noted a gradual accumulation of *SFRP2*+ fibroblasts from the pleural surface towards the Immunofibrotic Zone (**Fig. 3N)**. This is contrasted by the significantly lower densities of CD4+ T cells and *SFRP2*+ fibroblasts in the Elastofibrotic Zone, reflective of a post-remodeling state.

In direct contact with the underlying edge of alveolar-to-elastofibrotic remodeling, we identified *CTHRC1*+ fibrotic fibroblasts embedded in immature basophilic ECM (**Fig. 2H**, **S3**). Deposition of immature collagen, lack of elastic fibers in the EvG stain (**Fig. 2H**) and presence of fibrotic fibroblasts in this zone (**Fig. 3H, A**) are hallmark histopathological characteristics otherwise found in fibroblastic foci of IPF lungs^17,33,34^. Within this zone, the strong spatial expression of *COL1A1* was paralleled by IF detection of CTHRC1 (**Fig. 2K**), confirming *COL1A1*^hi^ cells as CTHRC1+ fibrotic fibroblasts. Histological quantification of *COL1A1*^hi^ cells expectedly demonstrated the highest densities in the Usual Fibrotic Niche. Some *COL1A1^hi^* cells exhibited co-expression of *SFRP2*. Topographically, these *SFRP2*+*COL1A1^hi^* cells were mainly restricted to the Immunofibrotic Zone and the Usual Fibrotic Niche (**Fig. 3N**). As *CXCL14* expression was also observed in mesenchymal cells located in the Usual Fibrotic Niche (**Fig. 3D**), *CXCL14* expressing *LEPR*+*ITGA8*+*DIO2*+ elastofibrotic fibroblasts may reflect this intermediate mesenchymal state that can differentiate in *CTHRC1*+ fibrotic fibroblasts. Accordingly, ssGSEA suggested a fibrotic activation state with induced collagen synthesis in elastofibrotic fibroblasts (**Fig. 2L**).

The Usual Fibrotic Niche was spatially restricted to a cuboidal-shaped layer of Aberrant Basaloid cells continuously overlined by *CTHRC1*+ fibrotic fibroblasts, resulting in a sharp demarcation line between the remodeled fibrotic tissue and the alveolar parenchyma (**Fig. 4H**). This zone will be further explored in the next paragraph.

**Figure 4.**
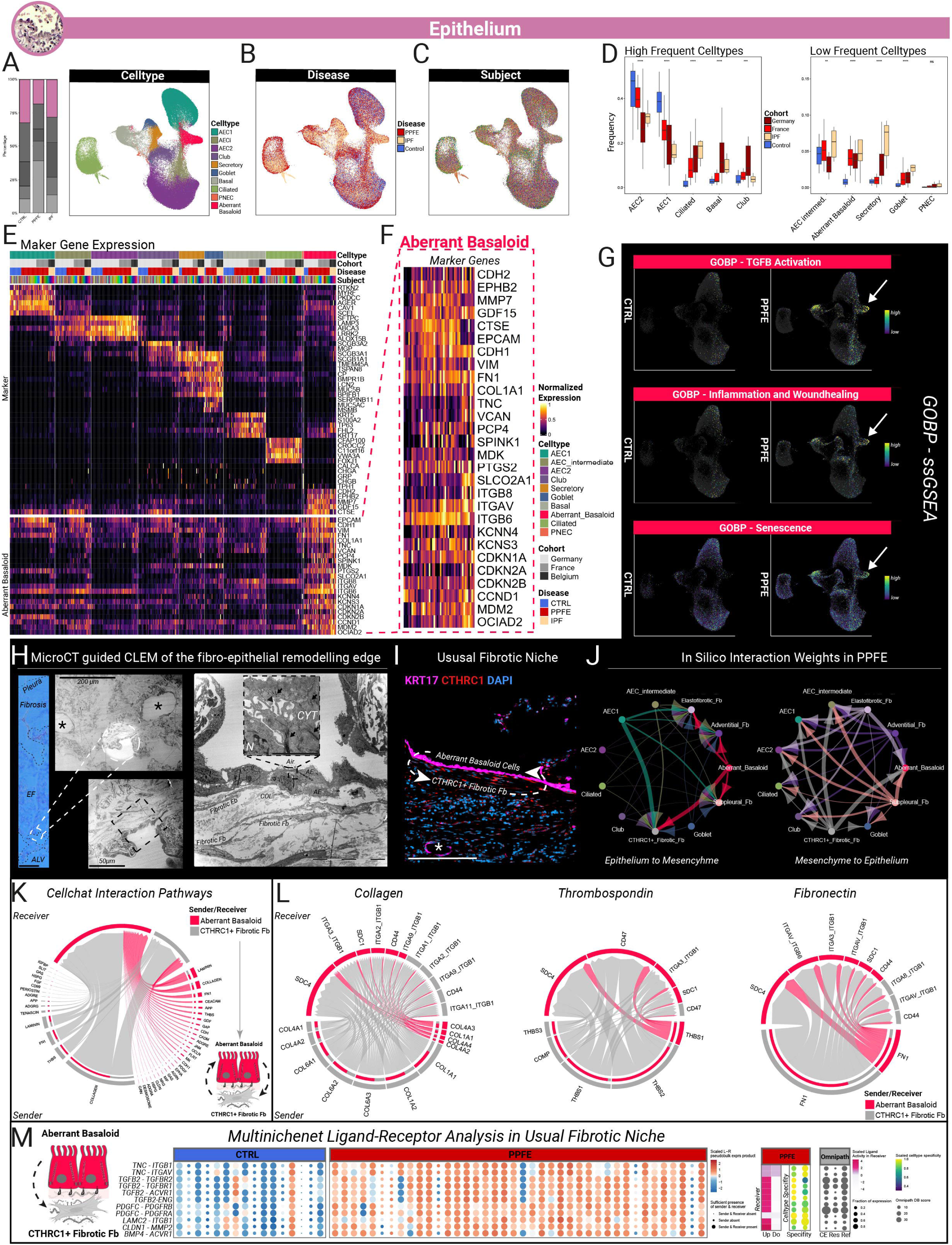
Sub analysis of the epithelial cell lineage. **A** Relative fraction of epithelial nuclei and UMAP embedding of identified epithelial cell types (AEC1, AECintermediate, AEC2, Club, Secretory, Goblet, Basal, Aberrant Basaloid, Ciliated and PNEC) within the epithelial lineage **B** UMAP embedding by disease. **C** UMAP embedding by disease **D** Boxplots display the relative distribution of each epithelial cell type relative to the total epithelial lineage compartment, stratified by disease-related cohort. Whiskers indicate 1.5 times the interquartile range (IQR). Statistical comparisons between PPFE-GER, PPFE-FR, CTRL and IPF were performed using Kruskal-Wallis test with *: p ≤ 0.05, **: p ≤ 0.01, ***: p ≤ 0.001, and ****: p ≤ 0.0001. Post-hoc tests were performed under use of the Wilcoxon Rank Sum test, with Bonferroni correction for multiple comparisons across cohorts and celltypes. Adjusted p-values less than 0.05 were considered statistically significant. Detailed statistics are provided in supplemental table E8. **E** Unity scaled average expression of canonical markers per subject (upper heatmap) and extended aberrant basaloid marker genes (lower heatmap). Subjects were grouped after cohort and disease. Each subject was assigned to a unique color. **G** Featureplots of cell based single sample gene set enrichment analysis (ssGSEA) enrichment values of distinct gene ontology biological pathways (GOBP) involved in fibrotic remodeling split by disease and plotted for CTRL and PPFE. Arrows indicate aberrant basaloid cells. **H** Micro-CT guided correlative imaging of early septal thickening beyond the fibro-alveolar remodeling edge. Low resolution image of OsO4 contrasted 50µm thick sections (left panel, scale bar is 500 µm). Pleura and border between subpleural fibrosis and AFE remodeling are indicated by a dashed line. Scale bars resemble 50µm (low magnification), 20 µm (medium magnification) and 10 µm (high magnification). Dashed boxes indicated areas of interest for higher magnification imaging. Dashed boxes resemble areas of interest. Nuclei (N) and cytoplasm (CYT) are annotated on high magnification images. Arrows indicate intra-cellular contrasted intermediate filaments. **I** IF stains of the Usual Fibrotic niche and its overlining epithelium. **J** Inter-cellular ligand-receptor interaction weight analysis among major epithelial and mesenchymal celltypes split by assigned sender lineage. Edge thickness resembles interaction weight. Nodes are colored corresponding to assigned celltype colors in A except CTHRC1+ fibrotic fibroblasts were colored in grey for improved visual clarity. Interaction weight analysis was conducted with the R package CellChat v2. **K** Inter-cellular pathway-based interaction analysis between CTHRC1+ fibrotic fibroblasts and Aberrant Basasloid cells. Chord thickness corresponds to the number of predicted significant interactions between sender and receiver related to the respective pathway pathway **L** Chord diagram of predicted intercellular ligand-receptor pairs between Aberrant Basaloid cells and CTHRC1+ fibrotic fibroblasts related to top three pathways predicted in J. **M** MultiNicheNetR differential ligand-receptor analysis between PPFE and CTRL focused on Aberrant Basaloid-to-CTHRC1+ fibrotic fibroblasts signaling. Heatmap displays z-score normalized pseudobulk products of ligand and receptor expression in Aberrant Basaloid cells and CTHRC1+ CTHRC1+ fibrotic fibroblasts, respectively (left panel). Scaled ligand activity is shown for PPFE and CTRL, respectively. Specifity of ligand-receptor interaction within all celltype-celltype combinations, curation effort (CE), number of found references (Ref) and number of available resources in the Ominpath database is provided as well (right panel). Illustrations were created with biorender.com.

### Aberrant basaloid cells line the fibro-epithelial remodeling edge

Aberrant Basaloid cells were identified based on the expression of 28 marker genes^10^, which were fully conserved in both PPFE and the IPF cohorts (**Fig. 4F**). ssGSEA confirmed previously described pathogenic features of these cells by enrichment of gene ontology (GO) pathways involved in fibrotic pulmonary remodeling (**Fig. 4G**, **table E9**). IF staining for KRT17, COX2, ITGAV/B6, and CTSE confirmed their localization as a cellular monolayer lining the edge of elastofibrotic remodeling (**Fig. 5A,D**). This spatial pattern was consistently observed in all PPFE samples (**Fig. 4I**, **Fig. 3A,I**). Micro-CT guided ultrastructural imaging of the Usual Fibrotic Niche visualized Aberrant Basaloid cells as cuboidal shaped cells with apical microvilli and numerous cytoplasmic intermediate filaments (**Fig. 4H**, arrows), as already reported in IPF^35^. Within the Elastofibrotic Zone, we observed lesional remnants of alveoli and duct-like structures (*, **Fig. 4H,I**) lined by transitional epithelial cells co-expressing AEC2 (SFTPC, LAMP3), AEC1 (AGER), basaloid (KRT17) and injury markers (CTSE, **Fig. 5D**,**H**,**K**,**M**, **Fig. S5**). These intermediate cells were frequently observed in alveoli adjacent to the fibroelastotic remodeling edge as well (**Fig. 5F, G, Fig. S5**).

**Figure 5.**
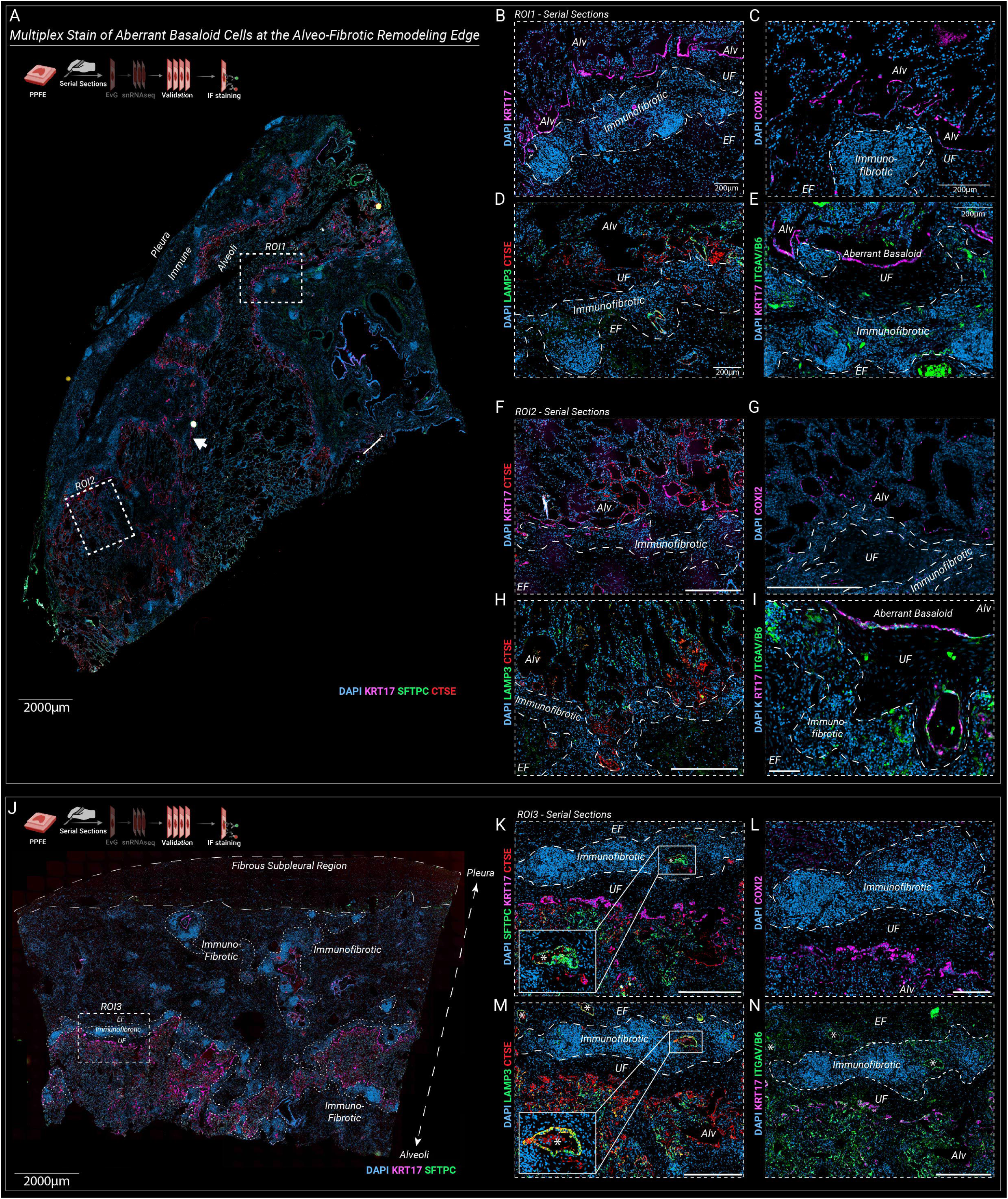
Immunofluorescence staining of Aberrant Basaloid cells in the PPFE lung. **A** Low magnification image of whole slide IF stained PPFE specimen. Dashed boxes indicate regions of interest. The arrow indicates the fibroepithelial remodeling edge. **D-I** High magnification of Aberrant Basaloid marker genes on serial sectioned tissue specimen. The Immunofibrotic Zone is highlighted by a dashed line. **J** Low magnification image of whole slide IF stained PPFE specimen with pronounced subpleural fibrosis (coarse dashed line). Dashed boxes indicate regions of interest. Immunofibrotic Zone is highlighted es well (fine dashed line). **K-N** High magnification of Aberrant Basaloid marker genes on serial sectioned tissue specimen. The Immunofibrotic Zone is highlighted by a dashed line. An intra-lesional remnant of alveoli (*) formed by epithelial cells being co-positive for LAMP3 and CTSE is shown with high magnification. Elastofibrosis Zone (EF), Usual Fibrotic Niche (UF), and alveoli (Alv) are annotated throughout IF stains. Scale bars are 200µm unless stated otherwise.

CellChat based interaction weight analysis highlighted *CTHRC1*+ fibrotic fibroblasts as a general signaling hub with highest connectivity to aberrant basaloid cells (**Fig. 4J**). Major involved pathways in this unsupervised inter-cellular crosstalk analysis were related to secretion of collagen, laminin, thrombospondin and fibronectin (**Fig. 4L)**. Aberrant basaloid cells express various stimulatory ligands such as *TGFB2* and *PDGFC*, with predicted activation of *CTHRC1*+ fibrotic fibroblasts in their surrounding niche via corresponding receptors (*TGFBR2*, *PDGFRB*, *PDGFRA* and *ACVR1*, **Fig. 4M**).

### Multimodal profiling of lesional vascular conglomerates in PPFE

Distribution of intra-endothelial cell frequencies was shifted in the PPFE group towards two ectopic cell populations (**Fig. 6D-F**): First, lymphatic ECs (*PDPN*, *CCL21*, *RELN*) were increased in PPFE (adj.pval_vs.CTRL_<0.001), but not in IPF (**Fig. 6E-F**). IF staining of PDPN revealed an expansion of lymphatic vasculature in AFE lesions (**Fig. 6G-I**). Overlay with phase contrast microscopic images located lymphatic EC in the same niche as elastic fiber depositing adventitial fibroblasts (**Fig. 6G**). Second, we observed a relative increase in ectopic systemic-venous endothelial cells (*COL15A1*, *PLVAP*, **Fig. 6A-F**). This endothelial phenotype had recently been described in IPF lungs^10,36^, non-specific interstitial pneumonia^37^ and RAS^38^. Co-localization of PPFE-associated endothelial and mesenchymal populations prompted us to (**Fig. 6G**) assess the underlying inter-cellular signaling circuits (**Fig. 6K-L**). Lymphatic endothelial cells had the highest scaled ligand activity for the *BMP2*-*ACVR1* and *PDGFC*-*PDGFRA* axis for signaling towards PPFE-associated fibroblasts (**Fig. 6L**), while systemic-venous EC signalled via *JAM2-JAM3*, *JAM2-ITGB1* and *JAM2-ITGAV* (**Fig. 6L**). Downstream target genes in PPFE-associated fibroblasts were enriched in proteoglycan metabolism, angiogenesis, and ECM remodeling pathways (**Fig. 6M**, **table E25**).

**Figure 6.**
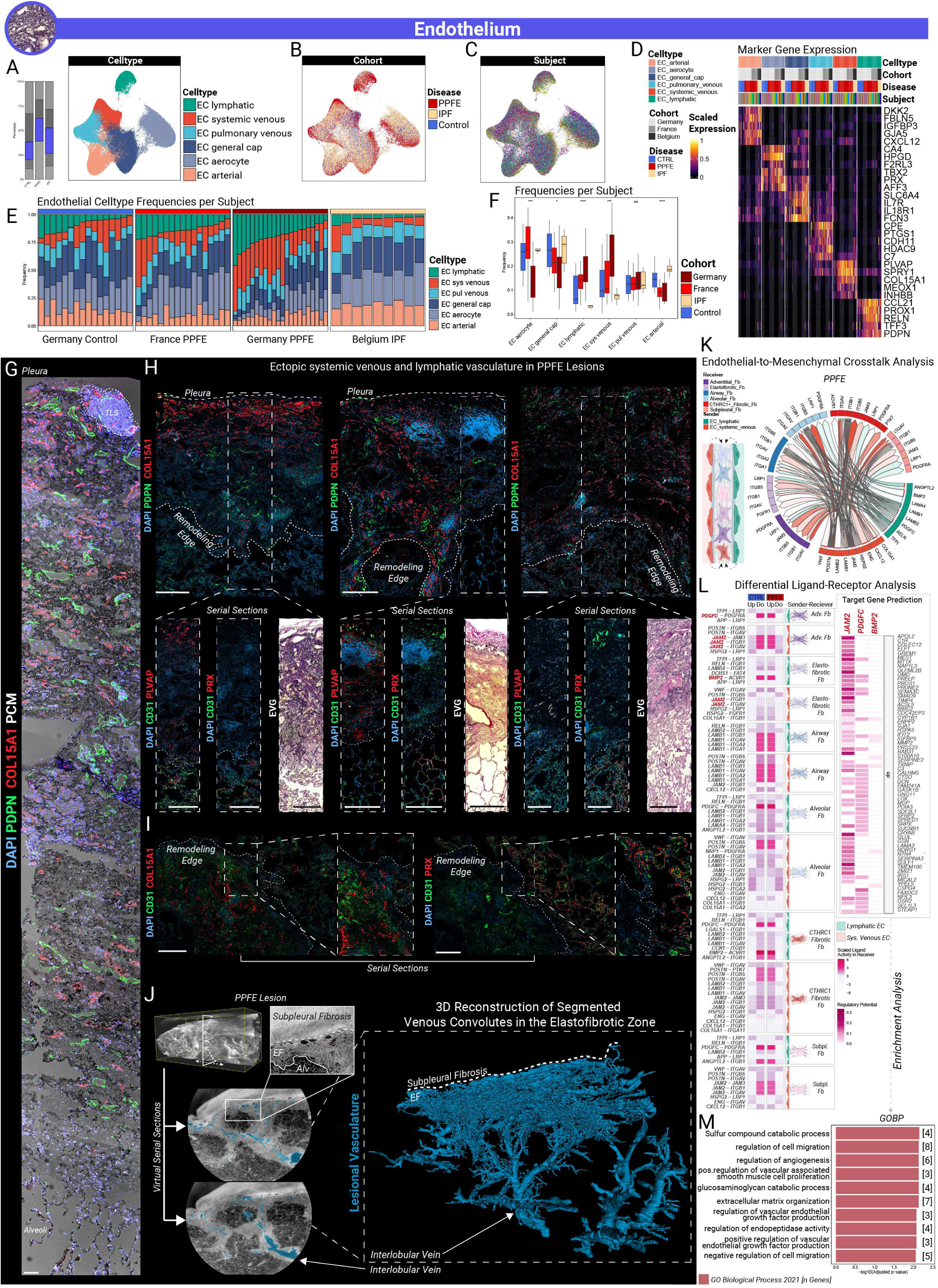
Ectopic endothelial cells in PPFE. **A** Relative fraction of endothelial nuclei in the global dataset and UMAP embedding of identified endothelial cell types (Lymphatic EC, systemic venous EC, pulmonary venous EC, general capillary EC, aerocytes and arterial EC) within the endothelial lineage **B** UMAP embedding by disease. **C** UMAP embedding by subject. **D** Unity scaled average expression of canonical marker genes per subject. Subjects were grouped after cohort and disease. Each subject was assigned to a unique color. **E** Stacked bar plots of celltype frequencies per subject colored by celltype and split by cohort. Subjects were ordered after frequencies of lymphatic EC. **F** Boxplots display the relative distribution of each endothelial cell type relative to the total endothelial lineage compartment, stratified by disease-related cohort. Whiskers indicate 1.5 times the interquartile range (IQR). Statistical comparisons between PPFE-GER, PPFE-FR, CTRL and IPF were performed using Kruskal-Wallis test with *: p ≤ 0.05, **: p ≤ 0.01, ***: p ≤ 0.001, and ****: p ≤ 0.0001. Post-hoc tests were performed under use of the Wilcoxon Rank Sum test, with Bonferroni correction for multiple comparisons across cohorts and celltypes. Adjusted p-values less than 0.05 were considered statistically significant. Detailed statistics are provided in supplemental table E18. **G** Overlay of phase-contrast light microscopy and IF staining for PDPN and COL15A1 across all remodeling zones from pleura to alveolar spaces. A tertiary lymphoid structure is encircled by a dashed line adjacent to the pleura. Scale bar is 200 µm. **H** IF staining of vascular markers in serial sectioned subpleural PPFE lesions for lymphatic EC (PDPN), systemic venous EC (COL15A1, PLVAP) and aerocytes (PRX). Scale bars represent 200µm unless stated otherwise. Pleura and remodeling edge are annotated by dashed lines. High magnification images were complemented with serial sectioned *EvG* stains. **I** IF staining of vascular markers in serial sectioned PPFE lesions related the interlobular septum for lymphatic EC (PDPN), systemic venous EC (COL15A1) and alveolar capillaries (PRX). Scale bars represent 200µm. The remodeling edge is annotated by dashed lines. High magnification images were complemented with serial sectioned *EvG* stains. **J** Zoomed hierarchical phasecontrast CT imaging (HiP-CT) of PPFE lesions located in the upper lobe. Pulmonary vein segmentation was performed employing an adopted nnU-Net framework. The resulting segmentation of the ectopic vasculature (blue). To enhance visual clarity, the 3D visualization shows the main vessel structures from the highlighted region illustrating an interconnection of interlobular veins and venous convolutes at the border of subpleural and elastofibrotic remodeling. **K** Differential up-scaled ligand-interactions receptor (PPFE vs. CTRL) between systemic venous EC and lymphatic EC (sender) and diverse mesenchymal cell types (receiver) were predicted with MultiNicheNetR (top panel). **L** Heatmap of the ligand-receptor interactome of systemic venous and lymphatic ECs with inflammatory mesenchymal celltypes. Downstream expressed target genes of ligands with highest predicted activity (*PDGFC*, *BMP2* and *JAM2*) were inferred in adventitial and elastofibrotic fibroblasts (right panel). **M** Enrichment analysis for GOBP terms of predicted target genes. Detailed enrichment results are provided in table E25. Illustrations were created with biorender.com.

To obtain a better understanding of the three-dimensional context of the vasculature in PPFE lesions, we performed hierarchical phase-contrast tomography (HiP-CT) of the intralesional vasculature (**Fig. 6J**). 3D reconstruction indicates that the pronounced vascular conglomerate located at the border of the elastofibrotic and Subpleural Fibrosis Zone communicates with interlobar veins (**Fig. 6J**, **Supp. Video 1**), suggesting inter-vascular anastomosis between systemic-venous and pulmonary-venous vasculature. Micro-CT imaging of osmium tetroxide and phosphotungstic acid contrasted fixed lung tissue confirmed numerous small vessels infiltrating the remodeled PPFE tissue originating from the alveolar parenchyma via the remodeling edge (**Fig. S2A-B,** white arrows). Simultaneously, we noted a subpleural vascular offspring of narrow caliber vessels with centripetal rejuvenation (**S2A-B**, black arrows).

### Immune cell landscape in PPFE

Relative cell frequency analysis per subject among cohorts revealed a significant relative increase in the lymphoid cell proportions against CTRL and IPF, that was consistent among both PPFE cohorts (PPFE-GER: 44.6%±13.6%, PPFE-FR: 28.3%±13.6%, CTRL: 8.52%±13.76%, IPF: 12.6%±7.6%, adj.pval_vs.CTRL_<0.01 and <0.01; adj.p-val_vs.IPF_<0.01 and 0.02, **Fig. 1C-E**, **Table E4-5**). Surface fractions of remodeled tissue per sample positively correlated with proportion of lymphocytes (**Fig. S7,** Pearson’s R=0.43, pval<0.001). Phenotyping of lymphoid cells exhibited a broad repertoire of immune cells to emerge in the PPFE lung, further explored in the supplemental results. In contrast, lymphoid cell frequencies were not increased in IPF (**Fig. 1H**, adj.pval_vs.CTRL_=0.75, **table E8**).

Myeloid immune cell populations exhibited a relative decrease of alveolar macrophages (AM, PPFE-GER: 1.83±2.69%, adj.pval_vs.CTRL_<0.001; PPFE-FR:1.57±0.96%, adj.pval_vs.CTRL_<0.001, **Fig. 7E-G**). Interestingly, the AM frequency was unaltered in the IPF dataset, in which 22.56% of myeloid cells were identified as AM (adj.pval_vs.CTRL_=1). Further differences in cell frequencies between PPFE and IPF comprised increased proportions of classical monocytes within the lung tissue in PPFE (cMono, adj.pval_vs.CTRL_ each < 0.001). Remarkably, fractions of *SPP1*+*CHIT1*+ profibrotic macrophages were not elevated in the PPFE lung (adj.pval_vs.CTRL_=1).

**Figure 7.**
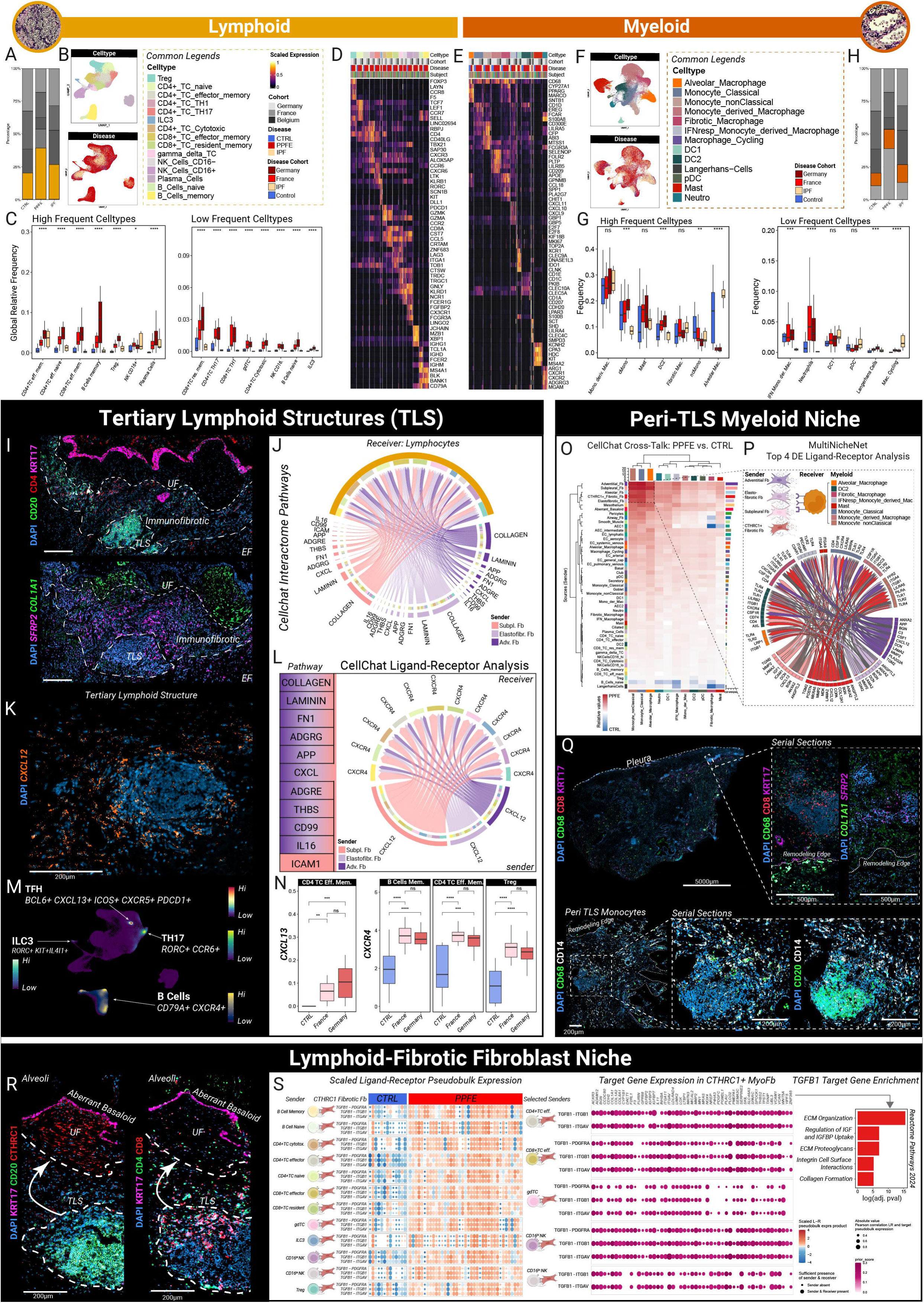
Diverse immune cell repertoire in PPFE. **A** Relative fraction of lymphoid nuclei in the global dataset. **B** UMAP embedding of identified lymphoid cell types (Tregs, CD4+ T cells (TC) naïve, CD4+ TC effector memory, CD4+TC TH1, CD4+ TC TH17, ILC3, CD4+ TC cytotoxic, CD8+ TC effector memory, CD8+ TC resident memory, gamma delta T-cells (gdTC), NK Cells CD16−, NK Cells CD16+, Plasma cells, B cells naïve and B cells memory) within the lymphoid lineage (top panel) and UMAP by disease (bottom panel) **C** Boxplots display the relative distribution of each lymphoid cell type relative to all nuclei from all lineages per sample, stratified by disease-related cohort. Whiskers indicate 1.5 times the interquartile range (IQR). Statistical comparisons between PPFE-GER, PPFE-FR, CTRL and IPF were performed using Kruskal-Wallis test with *: p ≤ 0.05, **: p ≤ 0.01, ***: p ≤ 0.001, and ****: p ≤ 0.0001. Post-hoc tests were performed under use of the Wilcoxon Rank Sum test, with Bonferroni correction for multiple comparisons across cohorts and celltypes. Adjusted p-values less than 0.05 were considered statistically significant. Detailed statistics are provided in supplemental table E18. **D-E** Unity scaled average expression of canonical marker genes of lymphoid and myeloid celltypes per subject. Subjects were grouped after cohort and disease. Each subject was assigned to a unique color. **F** UMAP embedding of identified lymphoid cell types (Alveolar Macrophages, classical monocytes (cMono), non-classical monocytes (ncMono), monocyte derived macrophages (Mono. Deriv. Mac.), fibrotic macrophages, interferon gamma response macrophages (IFN. Mono. Deriv. Mac.), cycling macrophages, dendritic cells type 1 (DC1) and type 2(DC2), Langerhans cells, plasmacytoid dendritic cells (pDC), mast cells and neutrophils) within the myeloid lineage (top panel) and UMAP by disease (bottom panel). **G** Boxplots display the relative distribution of each myeloid cell type relative to the total myeloid lineage compartment, stratified by disease-related cohort. Whiskers indicate 1.5 times the interquartile range (IQR). Statistical comparisons between PPFE-GER, PPFE-FR, CTRL and IPF were performed using Kruskal-Wallis test with *: p ≤ 0.05, **: p ≤ 0.01, ***: p ≤ 0.001, and ****: p ≤ 0.0001. Post-hoc tests were performed under use of the Wilcoxon Rank Sum test, with Bonferroni correction for multiple comparisons across cohorts and celltypes. Adjusted p-values less than 0.05 were considered statistically significant. Detailed statistics are provided in supplemental table E22. **H** Relative fraction of myeloid nuclei in the global snRNAseq dataset. **I** Tertiary lymphoid structure niche (TLS). Serial IF and ISH staining of colocalizing immune cells and fibroblasts. Dashed box indicates area of interest. **J** Inter-cellular pathway-based interaction analysis between subpleural fibroblasts, adventitial fibroblasts, elastofibrotic fibroblasts (sender) and lymphoid cell subsets (receiver). **K** ISH of *CXCL12* expression in the peri-TLS niche. Scale bar is 200µm. **L** Predicted interaction of subpleural fibroblasts, adventitial fibroblasts, elastofibrotic fibroblasts (sender) and lymphoid cell subsets (receiver). CXCL signaling as non-ECM related pathway is highlighted by depiction of the corresponding ligand receptor chord diagram. Chord thickness corresponds to CellChat v2 calculated interaction strength for each ligand-receptor pair per cell type pair. **M** Nebulosa plots of TLS formation related celltypes (ILC3, follicular T-cells, TH17 and *CXCR4*+ B cells). Displayed densities depend on co-expression of the referred marker genes. Nebulosa plots were plotted separately and stitched together for visualization purposes. Separate nebulosa plots are provided in Fig. S8. **N** Average per subject expression stratified by cohort of TLS formation related genes. Statistical comparison among cohorts was performed under use of Wilcoxon rank sum test with *: p ≤ 0.05, **: p ≤ 0.01, ***: p ≤ 0.001, and ****: p ≤ 0.0001. **O** Peri-TLS Niche. Comparative relative crosstalk weight analysis between PPFE and CTRL towards myeloid cells. The cluster of top signaling celltypes is highlighted by a dashed box. **P** Chord-diagram of differential ligand receptor analysis of top sending celltypes and myeloid cells employing MultiNicheNetR. **Q** IF staining of CD14 and CD68 expressing monocytes across remodeling niches in PPFE. Serial sections indicate co-localization of *SFRP2*+fibroblasts and CD8+T-cells with CD14+ monocytes. Dashed lines indicate zoomed in ROIs. **R** IF imaging of the TLS in proximity to the Usual Fibrotic niche (UF). Scale bar is 200µm. TLS and edge of the Usual Fibrotic niche are indicated by a dashed line. **S** MultiNicheNetR differential ligand-receptor analysis between PPFE and CTRL focused on Lymphoid-to-*CTHRC1*+ myofibroblasts signaling. Heatmap displays z-score normalized pseudobulk products of ligand and receptor expression in lymphoid cells and CTHRC1+ fibrotic fibroblasts, respectively (left panel). Predicted target gene regulation in signal receiving CTHRC1+ fibrotic fibroblasts in the means of Pearson correlated gene expression with pseudobulk products (>0.3). Color scale indicates prior knowledge-based regulatory potential (right panel). Enrichment of target genes was performed for Reactome Pathways 2024 terms. Illustrations were created with biorender.com.

### Immuno-mesenchymal cell communication networks partition into three distinct niches

Correlation of *in silico* cellular crosstalk analysis with *in-situ* mapping of lymphoid and myeloid cell types indicated three inter-cellular signaling hubs in PPFE lesions:

CD14+ monocytes that accumulated in the circumference of TLS exhibited an increased crosstalk weight from diverse fibroblast subsets as well (Peri-TLS Myeloid Niche, **Fig. 7O-Q**). Ligand-receptor analysis of the peri-TLS myeloid niche exhibited increased signaling weights between myeloid cells and adventitial, elastofibrotic, subpleural, and *CTHRC1*+ fibrotic fibroblasts (**Fig. 7O**). Of special interest is the fibroblast-to-monocyte communication through *CSF1-CSFR,* central for recruitment of classical and non-classical monocytes (**Fig. 7O,P**). IF staining localized aggregates of large CD68+ macrophages within alveoli at the remodeling edge (**Fig. 7Q)**. Smaller CD14+CD68+ monocytes were ubiquitously abundant in the Elastofibrotic Zone, while forming clusters in the periphery of TLS near surrounding CD8+ T-cells (**Fig. 7Q**). Serial sections for ISH indicated peri-TLS monocytes to assemble in the same topographic region as *SFRP2*+ fibroblasts, again suggesting an additive role for the predicted fibro-myeloid crosstalk in TLS formation of PPFE.

We observed an enhanced signaling circuit centered around inflammatory fibroblast populations and lymphocytes forming TLS (TLS Niche, **Fig.7H-N**). Within the TLS niche, *SFRP2*+ adventitial and elastofibrotic fibroblasts co-localized (**Fig. 3J**, **Fig. 7I)** with CD4+, CD8+, and CD20+ positive lymphocytes (**Fig. 7H,I,K**, **Fig. 3N**). Ligand-receptor analyses highlights signaling between *CXCL12+* fibroblast and lymphocyte subsets expressing the corresponding receptor *CXCR4* as prominent non-ECM-related pathway (**Fig. 7J**). Expression of *CXCL12* was increased in fibroblasts and *CXCR4* in lymphocyte subpopulations, suggesting the *CXCL12*-*CXCR4* axis as a potential link between recruited immune cells and inflammatory fibroblasts in PPFE (**Fig. S7A,N**). ISH confirmed *CXCL12* expression within the TLS niche (**Fig. 3J**, **Fig. 7K**). Targeted exploration of TLS-formation related cell types identified a *BCL6*+*ICOS*+*PDCD1*+ and *CXCR5*+ population of follicular T-helper cells within the cluster of CD4+ effector memory T-cells with increased expression of *CXCL13* in both PPFE cohorts (**Fig. 7N, Fig. S8**). In addition, we observed potential *RORC* expressing lymphoid TLS inducer cell populations such as TH17 cells as well as a minor population of ILC3 (**Fig. 7M**, **table E18**). Particularly, TH17 cells were abundant in the PPFE lungs and only to a significantly lesser extent in IPF lungs (**Fig. 7C**, adj.pval_vs.IPF_<0.001).

Last, we found a profibrotic lympho-fibroblastic crosstalk in accordance with the frequently observed spatial proximity of TLS to the Usual Fibrotic Niche (**Fig. 7R,S**). *SFRP2*+ *COL1A1*+ co-positive fibroblasts are enriched within the Immunofibrotic Zone (**Fig. 3N**) and localize in close proximity to *CTHRC1*+ fibrotic fibroblasts. Targeted ligand-receptor analysis revealed an upregulation of classical *TGFB1* signaling consistently within some of the lymphoid cell subsets being increased in the PPFE cohorts (**Fig 7S**). Significantly enriched pathways included ECM remodeling and collagen synthesis. Taken together, these findings suggest that local immune cells may catalyze local profibrotic mesenchymal phenotype conversion and fibrotic remodeling via *TGFB1* signaling.

## Discussion

With this study, we provide a first multinational snRNAseq atlas on the cellular and structural landscape of PPFE. On the cellular level, we observed unexpected activation states and diversity of mesenchymal cells in PPFE lesions, characterized by expression profiles reflective of a division of labor in driving elastofibrotic remodeling. We identified *CTHRC1*+ fibrotic fibroblasts and Aberrant Basaloid cells in PPFE, which had been discovered as fibroblast foci-forming cells in IPF before. We observed a pronounced expansion of lymphocytes in a disease that is considered purely elastofibrotic so far. On the histological level, we describe a zonation of the PPFE lesion, constructed by the above-mentioned PPFE-associated cell types. Comparison with IPF revealed common principles of lung fibrosis, especially the emergence of the Usual Fibrotic niche in two clinically divergent progressive fibrotic ILDs. But it also highlights differences, in particular PPFE-enriched inflammatory phenotypes of all major fibroblast populations, intriguing TLS formation in close proximity to the fibrotic remodeling edge, as well as intercellular circuits that maintain the zonation of PPFE lesions.

Inflammatory fibroblast activation states have been consistently reported in murine bleomycin-induced lung fibrosis, but its role in human disease has been unclear so far^23–25^. In PPFE, we show that multiple non-alveolar fibroblasts acquire such inflammatory states. Together, these fibroblast subtypes normally reside in the loose connective tissue space of the pleura, adventitia and the inter-lobular septum. Of special interest, *CXCL12* and *CXCL14* expressing *LEPR*+*ITGA8*+*SFRP2*+*DIO2*+ elastofibrotic fibroblasts^28^ are expanded in the PPFE lung, matching the reported signature of inflammatory fibroblasts in the bleomycin model. Surprisingly, this phenotypic subset was barely detectable in IPF. These inflammatory elastofibrotic fibroblasts exhibit substantial marker overlap with *CTHRC1*+ fibrotic fibroblasts, as reported for mice^23–25,28^. In murine bleomycin-induced lung fibrosis, *Cxcl12*⁺ inflammatory fibroblasts originate from alveolar fibroblasts and progressively differentiate into *Cthrc1*⁺ fibrotic fibroblasts^25^. This contrasts with PPFE, where the cellular counterparts appear to rather originate from pulmonary loose connective tissue fibroblasts. Both *CXCL12*-expressing inflammatory adventitial and elastofibrotic fibroblasts are readily found in PPFE, but not in IPF, and are spatially associated with histopathological features such as septal elastosis and elastic fiber deposition. In amalgamate, our molecular profiling of the PPFE mesenchyme emphasizes the relevance of inflammatory fibroblast states for elastofibrotic remodeling in human disease.

The role of inflammation in lung fibrosis has remained controversial over decades. On the one hand, various autoimmune diseases can lead to fibrotic remodeling of the lung, with inflammatory changes usually preceding the development of definitive fibrosis^39–41^. On the other hand, fibrosis can cause inflammation, as fibrotic remodeling itself induces autoantigen shedding with consecutive autoantibody formation^42^. In addition, fibrotic ILD patients exhibit a dysregulated immune compartment with expansion of Tregs, GZM^high^ CD4+ and CD8+ effector T-cells^43,44^. However, untargeted immunosuppression increases mortality in IPF, especially in patients with short telomers^45,46^, which has also been observed in telomeropathy-associated PPFE^46–48^. Therefore, anti-inflammatory treatment regimen are rarely tested in fibrotic lung diseases. Our work highlights accumulating immune cells forming TLS and accompanying inflammatory fibroblasts as integral components of the cellular landscape of PPFE, providing a molecular rationale for niche-specific therapeutic strategies. This may apply beyond PPFE, as *SFRP2*+ fibroblasts have been described as myofibroblast precursors in systemic sclerosis skin lesions^49^, while *CXCL12+* fibroblasts shape the TLS-related stromal niche in various cancers^32^ and drive fibrotic bone marrow remodeling in primary myelofibrosis^50^. In the lung, inflammatory fibroblasts seem to serve as orchestrators of immune cell recruitment and TLS formation culminating in downstream formation of the Usual Fibrotic Niche. Hence, our work raises the question whether targeting inflammatory fibroblasts or TLS formation by interference with the central *CXCL12*-*CXCR4* axis could alleviate immunofibrotic remodeling in PPFE.

The divergent radiological and histopathological features of PPFE and IPF suggest different disease processes. It was correspondingly surprising for us that *CTHRC1*+ fibrotic fibroblasts and Aberrant Basaloid cells emerged in the PPFE lung as well. As both pathological cell types recapitulate hereby their inter-cellular framework in PPFE and IPF, we emphasize their spatial co-emergence as “Usual Fibrotic Niche”. Aberrant Basaloid cells are the key players in fibrotic remodeling due to their unique characteristics of TGFβ activation, senescence and injury memory^10^, while *CTHRC1*+ fibrotic fibroblasts secreted immature collagen^33,34^. Since their discovery a few years ago, an increasing number of studies also report Aberrant Basaloid cells in IPF and other fibrotic lung diseases^10,13,38,40^. In the IPF lung, *CTHRC1*+ fibrotic fibroblasts aggregate to fibroblast foci, which are covered by Aberrant Basaloid cells^10,13^. In contrast to IPF, the site of active remodeling of PPFE is not a patchy chaotic mosaic but a continuous layer at the alveolo-fibrotic remodeling edge (**Fig. 8C**). PPFE as disease highlights therefore that the Usual Fibrotic niche is more common than previously anticipated and not limited to the gestalt of fibroblastic foci.

**Figure 8.**
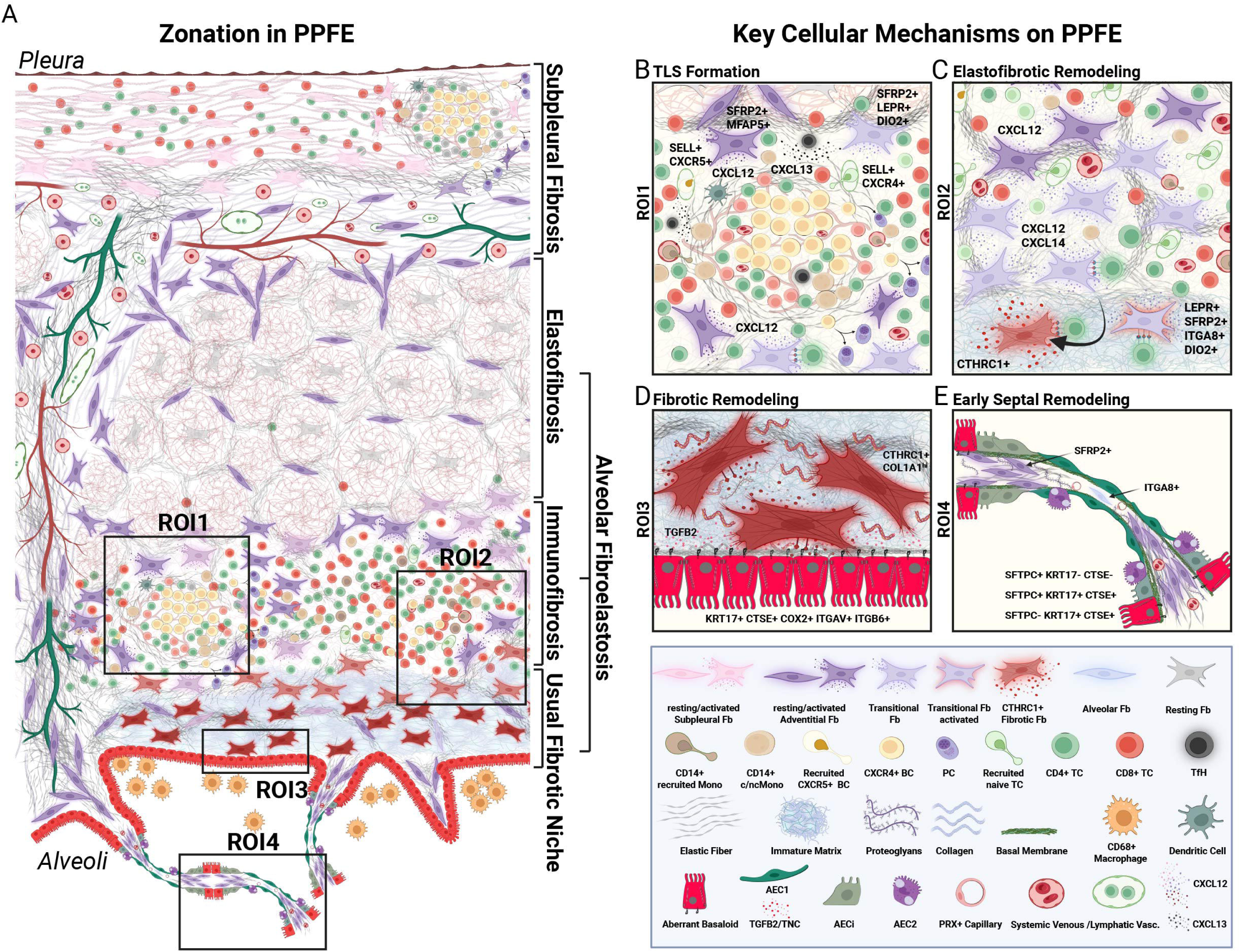
Niches of Remodeling in PPFE. **A** Schematic overview of distinct remodeling related niches in PPFE. **B**-**E** Zoomed in pathogenic niches with related key cellular mechanisms contributing to AFE remodeling.

As in this PPFE study, the loss of *CD31*+*PRX*+ capillaries and the emergence of ectopic *COL15A1*+ *PLVAP*+ vessels^36^ have been consistently reported in IPF^10^, non-specific interstitial pneumonia (NSIP)^37^, and RAS^38^. The extent and communication of ectopic *COL15A1*+ *PLVAP*+ vascular convolutes in the Elastofibrosis Zone with interlobular pulmonary veins in the 3D Hip-CT segmentation in the PPFE lung is somewhat surprising. In contrast to IPF, we observed an expansion of lymphatic ECs, localized to the subpleural and interlobular lymphatic vasculature. *CCL21*+*PDPN*+ lymphatic EC may enhance recruitment of *SELL* expressing naïve T cells to sites of ongoing elastofibrotic remodeling further shaping the micromilieu of the Immunofibrotic Zone. Together, our PPFE study supports that the observed alveolar vascular remodeling is a general phenomenon of fibrotic ILD, and elaborates on the unique contribution of the lymphatics to elastofibrotic remodeling.

In summary, this study represents an unprecedented cellular survey of PPFE, in which ectopic and aberrant cell populations spatially organize in a unique zonation pattern. We reveal PPFE-enriched inflammatory fibroblast activation states underlying the immunofibrotic remodeling that promote elastosis, and highlight Aberrant Basaloid cells and *CTHRC1*+ fibrotic fibroblasts as core constituents of the Usual Fibrotic Niche, driving collagenous fibrotic remodeling. The PPFE atlas is accompanied by an easily accessible webtool, and provides a molecular rationale for PPFE-targeting and ILD-overarching niche-specific therapeutic strategies.

## Supporting information

PPFE Supplements

## Acknowledgments

We are grateful to all patients and control subjects who participated in this study. Special thanks to Leon Giercke from the Schupp Lab at Hannover Medical school for his support with the IF and ISH imaging. We thank Susanne Fassbender and Susanne Kuhlmann of the institute of functional and applied anatomy for their great expertise in TEM sample preparation. We also want to express our gratitude to Regina Engelhardt and Christina Petzold-Mügge from the Department of Pathology at Hannover Medical School for their help and expertise in localizing and sectioning for the archived FFPE blocks. We are also thankful to the Fiedler and Braun research groups for generously providing laboratory equipment.

## Authors Contribution

JR, JCS, AC, AJ, LN and JCK assembled the PPFE cohorts and extracted clinical data. JG, MG, FI, KA, HM, and PM supervised the transplant program and provided clinical data. JCS, NK, BV and WW provided data on the IPF comparator cohort. SD and MPD reviewed the HRCTs of screened patients. AC, LN, VTM and DJ reviewed histology. AC, JCK and LN supervised the biobank and sample processing. JR and LK performed the micro-CT. MB performed flow sorting of nuclei. LML, JR, LC and JCS performed the snRNA experiments. JR, JCS and AJ performed the snRNAseq analysis. JR and LC performed the IF and RNA-ISH validation experiments incl. quantitative analysis. JR, JH and LK performed and analyzed correlative micro-CT and electron microscopy. AS, RE, DY, SEV and JJ performed HiP-CT imaging and analysis. SG and JF assembled the interactive webtool. All authors provided critical interpretation, review, and commentary on data and the manuscript. The manuscript was drafted by JR, AJ and JCS and reviewed and edited by all authors. JCS and AJ conceptualized and supervised the study, and acquired funding.

## Material and Methods

A detailed description of the methods is provided in the supplements.

