## Supplementary material for "The pleuroparenchymal fibroelastosis atlas reveals aberrant cell states and their zonation as an alternate roadmap to lung fibrosis": PPFE Supplements: PPFE_Manuscript_Supplement_prefinal_05072025.pdf

### Supplemental Results

#### Mesenchymal Cell Repertoire

Within fibroblasts, we identified resident alveolar fibroblasts (*ITGA8*, *FRAS1*), subpleural fibroblasts (*HAS1*, *WT1*), adventitial fibroblasts (*MFAP5*, *PI16*, *SCARA5*) and airway fibroblasts (*WNT5A*, *COL10A1*) based on expression of distinct marker genes (**Fig. 2A, table E15**). In addition, vascular related cells including pericytes (*COXI2*) and smooth muscle cells (*DES*, *MYH11*) were abundant throughout all cohorts.

#### Aberrant Basaloid Cell Features

Within the epithelial subset both PPFE cohorts we identified Aberrant Basaloid cells (*KRT17*, *SOX4*, *SOX9*) with expression of genes involved in TGF-β1 activation (*ITGAV*, *ITGB6*), senescence (*CDKN1A*, *CDKN2A*, *CCND2*, *MDM*) and epithelial-to-mesenchymal transition (*EMT*, *CDH2*, *COL1A1*, *VIM*). The frequencies of Aberrant basaloid cells were consistent through both cohorts, similar to the frequencies in IPF, and significantly increased compared to controls (PPFE-GER; 4.22±2.91%, $pval_{vs.Control}<0.001$ and PPFE-FR: 5.03±3.78% $pval_{vs.Control}<0.001$ (**Fig. 4A-E**), IPF 5.83±4.17%, $pval_{vs.Control}=1$, CTRL 1.01±1.01%). Aberrant Basaloid cells were found in each investigated PPFE and IPF sample (**table E10**).

#### Endothelial Cell Repertoire

Relative frequencies of endothelial cells (*CDLN5*+ *CDH5*+ *CLEC14A*+[1]) were reduced in both PPFE datasets compared to IPF (PPFE-GER: 7.89±4.19%, $pval_{vs.IPF}<0.001$; PPFE-FR: 10.16±4.19% $pval_{vs.IPF}<0.001$; **Fig. 1H**, **table E19**). Compared to the control frequencies, only the PPFE-GER cohort exhibited decreased frequencies of endothelial cells (PPFE-GER, $pval_{vs.Control}=0.003$; PPFE-FR, $pval_{vs.Control}=1$, **Fig. 1H**). All major cell types including arterial EC (*DKK2*, *FBLN5*), aerocytes (*PRX*, *HPGD*), general capillary EC (*FCN3*, *RGCC*), pulmonary venous EC (*PTGS1*, *CPE*)[2] were detected throughout all investigated cohorts (**Fig. 6A-C**).

#### Lymphoid Cell Repertoire

The highest frequencies observed in PPFE for CD4+ effector memory TC (PPFE-GER: 10.38±3.49%, PPFE-FR: 9.59±3.13%), naïve CD4+ TC (PPFE-GER: 11.07±3.10%, PPFE-FR: 12.33±3.66%), CD8+ effector memory TC (PPFE-GER: 11.31±4.25%, PPFE-FR: 12.00±4.90%) and memory B-Cells (PPFE-GER: 16.47±9.05%, PPFE-FR: 11.91±9.08, canonical marker genes are listed in **Fig. 7 D** , **table E23**). Comparative analysis of lymphoid cell type frequencies relative referenced to total number of nuclei per sample indicated significant and reproducible increases in PPFE for naïve B cells, naïve CD4+ TC, but also TH1 CD4+ TC, cytotoxic CD4+TC, CD8+ effector memory TC, gdTC and CD16$^{lo}$ NK cells in PPFE compared to CTRL and IPF (**table E21**, **Fig. 7C**, $pval_{vs.CTRL}<0.01$). Taking together, these findings highlight a major pro-inflammatory microenvironment in the PPFE lung marked by influx of CD4+ TC and B cells.

### Material and Methods

#### Patient and Sample selection

Diagnosis of PPFE was made in concordance with current guidelines in the literature on multidisciplinary team discussion[3]. CT pattern fulfilling the criteria definite or compatible PPFE were considered eligible for the study. Samples from patients that had undergone re-transplantation in the means of PPFE in the context of restrictive allograft syndrome were excluded from further analysis. In the time frame from 2015 – 2024 we retrospectively identified a total of 23 PPFE patients at a German Center and 17 patients at a French center. Biobanked formalin-fixed paraffin embedded biobank material from both cohorts was screened for the combination of pleural fibrosis and alveolar fibroelastosis (AFE). Screening involved micro-CT imaging (n=10) and *Elastica van Gieson* or Orcein stainings. In addition, lung tissue from downsizing lungs or peripheral tumor resections served as control (n=7). Data from additional 9 controls published previously by our group (GSE284081) were also used for comparative downstream analysis. 1 FFPE block per patient was selected for further processing. Written informed consent was obtained from all subjects. The study was approved by the local ethics committee of the Hannover Medical School of Hannover, Germany (Ethic vote #11032_BO_K_2023), APHP, Bichat Hospital (Comite de Protection des Personnes Ile de France 1, No. 0811760) and Marie Lannelongue Hospital (Ministère de l’Enseignement Supérieur de la Recherche et de l’Innovation –AC-2020-4284).

#### Nuclei isolation from FFPE specimens for Chromium Fixed RNA Profiling for Multiplexed Samples

Solutions and buffers were prepared as follows and placed on ice:

| **Xylene + ethanol immersions** | | | |
|---|---|---|---|
| **Reagent** | **Stock Conc.** | **Final Conc. (%)** | **Vol per sample** |
| Ethanol | 100 % | 100/70/50/30 | 3.5 mL |
| Nuclease-free H20 | - | - | 1.5 mL |
| Xylene | Min. 98.5 % | 100 | 3 mL |

| **1X PBS + 0.5 mM $CaCl_2$** | | | | |
|---|---|---|---|---|
| **Reagent** | **Stock Conc.** | **Final Conc** | **Vol/sample (3 mL)** | |
| PBS | 1X | ~ 1X | 2999.4 µL | |
| $CaCl_2$ | 2.5 M | 0.5 mM | 0.6 µL | |
| **Tissue Digestion Buffer (instead of Dissociation Solution) (2 mL/sample)** | | | | |
| **Reagent** | **Stock Conc.** | **Final Conc.** | **Vol for 2 mL** | **Vol for 32 mL** |
| PBS | 10X | 1X | 200 µL | 3200 µL |
| $CaCl_2$ | 2.5 M | 0.5 mM | 0.4 µL | 6.4 µL |
| Liberase TM | 2.5 mg/mL | 250 µg/mL | 200 µL | 3200 µL |
| Collagenase D | 100 mg/mL | 2.5 mg/mL | 50 µL | 800 µL |
| RNAse Inhibitor Protector | 40 U/ µL | 0.24 U/µL | 12 µL | 192 µL |
| Nuclease-free H20 | - | - | 1537.6 µL | 24601.6 µL |
| **Quenching Buffer (1 mL/sample)** | | | | |
| **Reagent** | **Stock Conc.** | **Final Conc.** | **Vol for 1 mL** | **Vol for 16 mL** |
| Nuclease-free H20 | - | - | 875 µL | 14 mL |
| Conc. Quench Buffer (10x Genomics PN 2000516) | 8X | 1X | 125 µL | 2 mL |

Three 50 µm-thick sections from each formalin-fixed paraffin-embedded (FFPE) tissue block were cut and transferred into a Miltenyi C-tube (Miltenyi #130-096-334). In a biological hood at ambient temperature, the specimens were deparaffinized and rehydrated in accordance with the following protocol: The removal of paraffin was achieved through the application of three 10-minute exposures to xylene. The volume of xylene utilized in each specimen and wash volume was at least 1 mL. In instances where 1 mL proved insufficient for complete coverage

of a specimen, additional xylene was incorporated. Rehydration was performed in sequential 1-minute series of 1-mL ethanol immersions: The mixture contained 200 milliliters of 100% ethanol, 100 milliliters of 70% ethanol, 50 milliliters of 50% ethanol, and 30 milliliters of 30% ethanol. Subsequently, each specimen was subjected to three washes with 1X Phosphate Buffered Saline (PBS) containing 0.5 millimeters of calcium chloride (2× 1 mL and 1× 800 mL or 3× 900 mL). Subsequent to the final wash, the maximum amount of excess liquid possible was extracted. In instances where feasible, a swinging-bucket rotor centrifuge, RNase/DNase-free wide-bore tips, and LoBind tubes were utilized. The Tissue Digestion Buffer (TDB) was subjected to a heating process at a temperature of 37°C for a duration of 10 minutes prior to its utilization. Two milliliters of TDB were added to each specimen, which was still in C-tubes. C-tubes were meticulously placed in a gentleMACS Octo Dissociator, and the following program was subsequently executed: 1. The first step in the procedure is to incubate the sample at 37°C for 45 minutes. This is followed by spinning the sample at 37°C for 30 seconds at a speed of (+)2,000 rpm (clockwise). The third step is to spin the sample at 37°C for 30 seconds at a speed of (-)2,000 rpm (counterclockwise). The final step is to detach the C-tubes. C-tubes were briefly (about 30 seconds) centrifuged at 300 rcf to collect all the cells/nuclei at the bottom of each C-tube. Each pellet was thoroughly resuspended in its supernatant.

The dissociated tissues were passed through 70 µm filters (PluriSelect #43-50070-51) to remove debris and undissociated tissue pieces. An additional wash of each C-tube and each 70 µm filter was performed. Therefore, 1 mL PBS was added to each C-tube, and the solutions were passed through the filters before rinsing each filter with 1 mL PBS. The filtrates were collected in the same tubes as the matching dissociated tissue. Then, specimens were centrifuged at 850 rcf for 5 minutes at 4 °C. Supernatants were removed without disturbing the tissue pellets. Each pellet was resuspended in 1 mL chilled Quenching buffer. Each specimen was passed through a 20 µm filter (PluriSelect #43-10020-50) to remove debris and undissociated cells. (If a very low nuclei count was expected, an additional wash of the filter was performed.) Afterwards, the nuclei concentration of each nuclei suspension was determined using a LUNA-FL™ Dual Fluorescence Cell Counter and Acridine Orange/Propidium Iodide (AO/PI) Cell Viability Kit: Each nuclei suspension was gently pipette-mixed. Then, 2 µL of AO/PI was mixed with 18 µL nuclei suspension. Disposable/reusable slides were filled with 10 µL of the AO/PI-Kit-sample-mix per chamber. Counting was done multiple times using fluorescence mode. Each nuclei suspension was stored as follows before proceeding with Flow-activated nuclei sorting (FANS): Enhancer (Chromium Next GEM RNA Profiling Sample Fixation Kit) was thawed for 10 min at 65 °C, then briefly vortexed and centrifuged. Enhancer was kept warm and the absence of any precipitate was ensured before use. 0.1 volume of pre-warmed Enhancer was added to each nuclei suspension and pipette-mixed. Samples were stored at 4 °C for up to 1 week.

#### Fluorescence Activated Nuclei Sorting (FANS) of DAPI+ single nuclei

The sorting of nuclei was conducted at the Cell Sorting Research Facility at Hannover Medical School. For the purpose of flow cytometric sorting, nuclei specimens were stained with 4′,6-diamidino-2-phenylindole (DAPI, Thermo Scientific™) immediately prior to sorting. The final concentration of DAPI ranged from 1 to 10 µg/mL, and was adjusted as needed to achieve optimal nuclei staining conditions. The sorting process was executed in the Central Research Facility Cell Sorting at MHH, employing cell sorters equipped with a 100 µm nozzle at a pressure of 35 psi (FACSAria III Fusion, FACSAria IIu, Becton-Dickinson, or MoFlo XDP, Beckman-Coulter). A total of 5 x $10^5$ DAPI+ single nuclei per specimen were sorted into individual 15-mL tubes, with each tube containing 1 mL of 0.5X PBS + 0.02% BSA. Intact single nuclei were isolated by flow cytometry based on light scattering properties and DNA content, measured through the fluorescence signal of the DAPI dye. A narrow gate was established in the DAPI channel to exclude nuclei with fractional DNA content and debris. In subsequent gating steps, doublets were eliminated using scatter pulse parameters, and the main population was refined further in an FSC-SSC gate. Reanalysis of 300 sorted nuclei from each specimen revealed a homogeneous population of nuclei. The sorted single nuclei suspensions were stored on ice and subsequently utilized for downstream analyses following the sorting of 16 specimens (Chromium Fixed RNA Profiling Reagent Kits for Multiplexed Samples).

#### 10x Genomics Chromium Fixed RNA Profiling for Multiplexed Samples

Sorted DAPI+ single nuclei were promptly processed further in accordance with the "User Guide, CG000527, Rev B, Chromium Fixed RNA Profiling Reagent Kits for Multiplexed Samples," which included the following choices and deviations: The multiplexing experiment was designed to "Maximize Number of Cells" without sub-pooling, and a post-hybridization pooled wash workflow was selected for all specimens. Hybridization took place in PCR tube strips. Given the initial sorting of 500,000 DAPI+ single nuclei per specimen and to minimize nuclei loss, the first counting step during post-hybridization was omitted. Furthermore, all specimens (16 specimens per pool) were completely pooled since the absolute number of nuclei was estimated to be nearly identical across all specimens due to FANS. Before passing the sample pool through a 20 µm filter (pluriStrainer Mini 20 µm, pluriSelect), the nuclei pellet was only resuspended in 25% of the volume indicated in the protocol to achieve a high nuclei concentration without needing to concentrate the nuclei suspension after counting (step “Cell Suspension Volume Calculator for Multiplexing 16 Samples”). A target of 128,000 nuclei was set for “Targeted Cell Recovery,” but a 20% larger volume of the nuclei suspension stock/reaction than specified was used based on the previously determined nuclei concentration, which was then brought up to 40 µL with Post-Hyb Resuspension Buffer. For the Sample Index PCR, a total of 12 cycles was deemed appropriate. Three different QC approaches were implemented per library after library construction: First, a 1:20 dilution of the stock library with Buffer EB (Buffer EB, Qiagen) was prepared, using 5 µL to determine a Qubit quantification of the diluted library. For this, Qubit™ 1X dsDNA Assay Kits (HS) + (BR) (Invitrogen™), Qubit™ assay tubes (Invitrogen™), and a Qubit™ 4 Fluorometer (Invitrogen™) were employed. Subsequently, 5 µL of each 1:20 diluted library were sent to the Research Core Unit Genomics (RCUG) at MHH for DNA fragment length analysis (HS) and Qubit quantification. Additionally, qPCR was conducted for each library using the KAPA Library Quantification Kit for Illumina® Platforms by Roche (KR0405 – v11.20), following their detailed protocol. The remaining stock libraries and 1:20 diluted libraries were stored at -20 °C.

#### RNA Library Quality Control

Following library construction, three distinct QC methodologies were executed. Initially, a 1:20 dilution of the stock library with Buffer EB (Qiagen) was prepared, utilizing 5 µL to ascertain the Qubit quantification of the diluted library. The Qubit™ 1X dsDNA Assay Kit (HS) + (BR) (Invitrogen™), Qubit™ assay tubes (Invitrogen™), and a Qubit™ 4 Fluorometer (Invitrogen™) were utilized in this process. Subsequently, 5 µL of each 1:20 diluted library were sent to the Research Core Unit Genomics (RCUG) at MHH for DNA fragment length analysis (HS) and Qubit quantification. Additionally, qPCR was conducted for each library using the KAPA Library Quantification Kit for Illumina® Platforms by Roche (KR0405 – v11.20), following their detailed protocol. The remaining stock libraries and 1:20 diluted libraries were stored at -20 °C.

#### Sequencing

cDNA libraries were subjected to sequencing on an Illumina NovaSeq platform at the Institute of Human Genetics (Hannover Medical School, Germany), employing a sequencing configuration of 28 base pairs for Read1, 90 base pairs for Read2, and 10 base pairs for Index5 and Index7. Subsequently, base calls were converted to reads utilizing Cell Ranger software (v7.1.0) and its bcl2fastq implementation, "mkfastq."

#### Data Processing

Subsequent processing of the reads was also performed using Cell Ranger (v7.1.0). The reads of the probes were aligned to the human reference genome GRCh38 (GENCODE v32/Ensembl 98, "GRCh38-2020-A") using the Chromium Human Transcriptome Probe Set _v1.0.1 for GRCh38-2020-A. Comprehensive technical details concerning the sequencing and data processing procedures can be found in the **Table E1** of the supplementary information. The count matrices were further analyzed using the Seurat pipeline (R package Seurat v 4.3.0.1) in R (v. 4.2.1, The R Foundation), following their recommendations for scRNAseq data processing and integration (https://satijalab.org/seurat/). UMI counts were normalized by applying a scale factor of 10,000 UMIs per cell. This was followed by natural log transformation with a pseudocount of 1 in accordance with methods previously published by our group and others [2,4,5].

#### Low-Quality filtering

Mitochondrial reads were identified by the "MT-" prefix to the gene identifier, and the proportion of mitochondrial reads was calculated for each nucleus. Low quality filtering was conducted in an iterative adaptive fashion per sample, following normalization, scaling, dimensionality reduction and plotting of lineage marker genes to control for information loss while filtering. Detailed information for the final applied filters for each sample regarding feature count, read count and percentage of mitochondrial reads can be found in **table E2** and **E3**. All nuclei with read counts < 100, feature count < 100 and > 15% mitochondrial were filtered out prior to downstream analysis. In preparation for the downstream integration of 23 German PPFE, 17 French PFE and 16 German CTRL datasets, pre-cleaned data objects of each sample were iteratively integrated and clustered using Seurat's recommended reciprocal principal component analysis (RPCA). Prior, Seurat's *FindVariableGenes* (default settings with nFeatures = 2000) function was used for feature reduction to identify the most variable genes for each subject. The patterns of common variance among these genes were used to integrate the datasets using *FindIntegrationAnchors*() and *IntegrateData*(), followed by scaling of the integrated expression matrix using *ScaleData*() as described previously[2].

#### Integration, Dimensionality Reduction und Clustering

The scaled values from the integration analysis were subjected to principal components selected based on their contribution to variance. These principal components were then employed to calculate Euclidean distances between nuclei in feature space, thereby creating a graph embedding in which nodes connected the nearest neighbors of each nucleus. Louvain cluster analysis was subsequently applied to this graph, and for visualization purposes, the distances of the nuclei and their graph embeddings were transformed using Uniform Manifold Approximation and Projection (UMAP). Clusters that did not exhibit significant differences in gene expression were merged. This iterative process was continued until all nuclei of all subjects were assigned to unique cell types whose characteristic features occurred consistently in all subjects. Subsequently, lineages (i.e. "Epithelium", "Mesenchyme", "Endothelium", "Lymphoid" and "Myeloid" cells) were defined by marker gene expression (**Fig. 1F**). This process was iterated minimum two times per cell lineage followed by graph embedding and respective cluster analysis to identify multiplets, empty cells and cellular debris, which were subsequently removed from the object prior to downstream analysis. Cleaned lineage datasets were subsequently integrated via RPCA integration followed by final dimensionality reduction and graph embedding as described above.

#### Clustering

Seurat's "*FindNeighbours()*" function was used with the default parameters of k = 20 and the Euclidean point distance as the metric for clustering the data set. Cluster resolution was increased until the algorithm failed to dissect major inter-cluster differential expression discrepancies.

#### Cell type Annotation

The determination of specific cell types was achieved by identifying cell type-specific marker genes using Wilcoxon rank sum tests under employment of Seurat's *FindMarkers* function, where all cells within a given cluster were compared to all other cells. To ensure the reliability of the results, all p-values were adjusted using Bonferroni's correction for multiple testing, and a p-value less than 0.05 was considered significant. To rank expressed marker genes for cell type identification, the diagnostic odds ratio (DOR) of each gene was calculated to better distinguish differentially expressed markers. Positive expression was defined as $X>0$. A pseudocount of 0.5 was used to avoid undefined values. The final annotation of cell types was performed in a supervised manner by comparing the DEGs (top 300 genes according to logDOR) of each cluster, as well as the expression of markers established in the literature. The final cell type annotation was performed by assigning the differentially expressed genes (DEGs) of each cluster as well as the expression of published cell markers[1,2,5–7].

#### Downstream integration with IPF snRNAseq 3' Chromium dataset.

A cleaned snRNAseq dataset from 9 IPF patients sampled at 3 distinct lung regions each (GSE286182)[8], was integrated with the cleaned PPFE and CTRL data to serve as disease control. Being mapped to a distinct reference genome (GENCODE_release37_GRCh38.p13), gene names were translated into their corresponding gene names from release version 32 under usage of the biomaRt suite and the respective gene annotation databases from ensembl (https://www.ensembl.org/, accessed latest 11/11/2024). Only features available in the 10X Flex assay were used for further downstream data analysis. Following further data preprocessing as indicated above, IPF snRNAseq data was sample wise integrated with PPFE and CTRL samples running Seurat's RPCA based reference-based integration considering samples derived from the main cohort as reference to avoid major batch effect due to distinct sets of features and chemistry. Samples per patient were pooled prior to statistical cell frequency analysis. Following integration, downstream dimensionality reduction, graphical embedding, clustering and celltype annotation was performed as described above.

#### Connectome and inter-cellular crosstalk analysis.

The R packages Cellchat (2.1.2) and MultiNicheNetR (v2.0.1) were used for (differential) Ligand-Receptor analysis[9–11]. Prior to each analysis the integrated snRNAseq dataset was randomly down sampled to up to 100 nuclei per sample and cell type. The down sampled dataset was subjected to the respective downstream pipeline under employment of recommended standard settings. For differential ligand-receptor analysis between PPFE and CTRL subsets we employed CellChatv2 and MultiNicheNetR, since the complementing nature of the underlying ligand-receptor databases of both packages.

For CellChatv2, snRNAseq objects for each condition (PPFE and CTRL) were separated and the count matrix of the RNA assay was subjected to the CellChat analysis pipeline ("*createCellChat()"*) employing default settings. All database categories, namely "Secreted Signaling", "ECM-Receptor", "Cell-Cell Contact" and "Non-protein Signaling" were incorporated into the downstream analysis. Communication patterns present in less than 10 cells were filtered out prior to further processing, employing more conservative "trimean calculation" [11]. Following identification of differentially expressed genes per cell type by Wilcoxon rank sum test (p-value = 0.05), inter-cellular communication probabilities were inferred by random walk-based network propagation technique. Likelihood of each ligand-receptor pair was aggregated on pathway level to compute pathway-based communication probabilities, which was conducted without prior normalization. Aggregated cell-communication networks indicating inter-cellular signaling weights were visualized as heatmap or circle or hierarchy plots, wherein edge thickness correlates with interaction weight or interaction counts, stored in distinct assays of the resultant CellChat object. Circosplots were used to visualize significant interaction pathways, which was tested before by permutation testing filtered for a p-value of 0.05[11].

For MultNicheNetR analysis the identical randomly subsampled subset of the snRNAseq dataset without prior condition-based separation was subjected to the default processing pipeline(https://github.com/saeyslab/multinichenetr/blob/main/vignettes/basic_analysis_steps_MISC.knit.md). MultiNicheNet ranks ligand-receptor interactions based on several factors: the differential expression of the ligand and its receptors, variations in ligand activity within the recipient cell type among samples, expression patterns specific to cell type and condition and the proportion of samples in the target condition that exhibit sufficient expression of the ligand-receptor interaction. Following pseudobulk based DE analysis under employment of the R package *edgeR* applied a quasi-likelihood negative binomial generalized linear model with empirical p-value estimation and false discovery rate (FDR) control via the Benjamini-Hochberg method to filter out significantly regulated ligands and receptors. Resulting pseudobulk expression libraries are normalized and log2 transformed. The results of the DE analysis are the basis of NicheNets ligand activity estimation, where the identification of upregulated and downregulated genes in receiver cell types is conducted per condition. The quantification of ligand activity was achieved through the implementation of area under the precision-recall curve (AUPRC), with default thresholds of logFC > 0.50 or < -0.50 and adjusted p-values < 0.05. To facilitate comparison across contrasts, Z-score normalization is applied to ligand activity values. Finally, target gene inference is conducted using the "*get_weighted_ligand_target_links*()" function from NicheNet, restricting targets to the top-ranked genes based on regulatory potential, applying a filter of a Pearsons's correlation of coefficient of 0.33. Final prioritization of ligands is conducted by weighted aggregation of logFC values for significant differentially expressed ligand and receptor genes, scaled ligand activity in receiver cells by min-max normalization, condition-and cell type specific log-transformed and normalized pseudobulk expression and fraction of samples per condition exceeding predefined threshold expression. The resulting list of ranked ligand-receptor pairs and their correlated expressed target genes in the respective receiver cell were visualized as circos plots, bubble plots or heatmaps. Scaled pseudobulk products, scaled ligand activity, cell type related expression specifity, correlated target gene expression and degree of interaction pathway curation in the "Omnipath" resource (https://r.omnipathdb.org/) were plotted separately the top ranked ligand-receptor pairings in the respective niche of interest. Downstream enrichment of Pearson correlated target genes in receiver celltypes was performed under use of the R package EnrichR (v3.2) for Reactome pathways (*Reactome_Pathways_2024*) or Gene Ontology Biological Pathways (*GO_Biological_Process_2021*). Bonferroni corrected adjusted p-values < 0.05 were considered statistically significant.

#### Pseudobulk Analysis

For pseudobulk analysis expression matrices were collapsed using Seurats *Aggregate_Epxression()* on subject level. Afterwards celltype/disease/subject instances were annotated. DESeq2[12] and its negative binomial distribution model were used for assessment of differentially expressed genes (DE genes) without prefiltering. Multiple testing was performed on the counts slot and was corrected via Bonferroni correction. Adjusted p-values < 0.05 were considered statistically significant.

#### Enrichment Analysis

For enrichment analysis significantly differentially expressed genes of selected mesenchymal subtypes were assessed on cell level employing MAST[13] testing. Genes with Bonferroni corrected p-values < 0.05 were considered statistically significant. Enrichment for either *Reactome Pathways* (v.1.82) or *Gene Ontology Biologic Pathways* (GOBP, v3.16.0) was performed with the R package ClusterProfiler v4.6.2. adjusted p-values < 0.05 were considered statistically significant. Graph based visualization of Enrichment terms was generated with the wrapper function *EnrichPlot* of the R package SCP (v0.5.6). This wrapper function utilizes the *Fruchtermann-Reingold* layout from igraph (v.1.4.0) for graphical representation of overlapped genes per enrichment term (edges) and number of enriched genes per term (nodes). Clustering of terms was performed under use of greedy clustering. For selected clusters of enrichment terms enriched gene-term interactions were represented as network plot using again an *Fruchtermann-Reingold* layout with edges connecting genes with their terms they enriched for.

#### single cell gene set enrichment analysis

Single samples gene set enrichment analysis (ssGSEA[14]) was performed with escape Package (v1.99.0) using default settings. ssGSEA on epithelial cells was used with the inbuilt GOBP database. Geneset for matrisome related genes was retrieved from the MatrisomeDB (https://sites.google.com/uic.edu/matrisome/matrisome-annotations/homo-sapiens) used for ssGSEA enrichment. Multiple testing was accounted for with Bonferroni Correction and pvalues < 0.05 were considered significant.

#### Immunohistochemistry (IHC)

IHC was conducted to evaluate the functionality and accuracy of primary antibodies. Prior to starting, all reagents were allowed to reach room temperature. A standard deparaffinization and rehydration procedure for FFPE tissue was carried out (2x 5 min xylene, 2x 3 min 100 % ethanol, 1x 3 min 95 % ethanol, 1x 1 min 70 % ethanol, 1x 1 min 50 % ethanol, 1x 5 min distilled water), followed by heat-induced antigen retrieval. For this, tissue slides were heated for 20 min at 98 °C in a 1× tris-based Antigen Unmasking Solution (Vector Laboratories, USA), which was diluted 1:100 with distilled water. After allowing the specimens to cool for 20 min, they were washed in 1x PBS for 5 min. Tissue sections were circled with a Roth®-Liquid Barrier Marker and placed in a wet chamber used to store slides during all subsequent incubation steps. Specimens were incubated for 10 min in BLOXALL Endogenous Peroxidase and Alkaline Phosphatase Blocking Solution (Vector Laboratories, USA) to inhibit endogenous peroxidase and alkaline phosphatase activity. Tissue slides were then washed for 5 min in 1x PBS and blocked for 20 min using a 2.5 % Normal Horse Serum Blocking Solution (Vector Laboratories, USA). Each specimen was incubated for 30 min with a primary antibody (diluted appropriately in 2.5 % Normal Horse Serum Blocking Solution) at room temperature. Following a wash of the tissue slides in 1x PBS for 5 min, each specimen was incubated for 30 min with a corresponding secondary antibody (anti-mouse, anti-rabbit, or anti-goat ImmPRESS reagent, conjugated with horseradish peroxidase, all Vector Laboratories, USA) at room temperature. The slides were washed again for 5 min in 1x PBS, and then incubated for 6-10 min in the ImmPACT® DAB Substrate Kit, Peroxidase (HRP) (Vector Laboratories, USA). The incubation time varied due to individual reaction times. After one final wash for 5 min in 1x PBS, specimens were counterstained in Hematoxylin Solution Gill no. 1 (Sigma-Aldrich, USA) for 3 min and then washed for 15 min in tap water (with intermittent exchanges of tap water). Subsequently, a standard dehydration in ethanol/xylene was performed (1x 15 sec 50 % ethanol, 1x 15 sec 70 % ethanol, 1x 1 min 95 % ethanol, 2x 1 min 100 % ethanol, 2x 3 min xylene). Tissue slides were mounted with VectaMount permanent mounting solution (Vector Laboratories, USA), and coverslips were placed on top. Stained slides were digitized and analyzed using a ZEISS Axioscan 7 and the ZEN 3.5 (blue edition) software. The slides were stored at room temperature.

#### Immunhistofluorescent (IF) stainings

Before starting, all reagents were allowed to equilibrate to room temperature. A standard deparaffinization and rehydration of FFPE tissue was carried out (2x 5 min xylene, 2x 3 min 100 % ethanol, 1x 3 min 95 % ethanol, 1x 1 min 70 % ethanol, 1x 1 min 50 % ethanol, 1x 5 min distilled water), followed by heat-induced antigen retrieval. For this, FFPE tissue slides were heated for 20 min at 98 °C in 1× tris-based Antigen Unmasking Solution (Vector Laboratories, USA), which was diluted 1:100 with distilled water. Afterward, specimens were cooled for 20 min in 1X PBS at room temperature. Meanwhile, a stock solution of 0.9995 X PBS + 0.05 % Tween® 20 was prepared to create 2.5 % normal serum (donkey/goat/horse) corresponding to the species of the secondary antibodies. All primary and secondary antibodies were diluted in the appropriate 2.5 % normal serum and stored at room temperature under aluminum foil until required. Once the slides had cooled, they were circled with a Roth®-Liquid Barrier Marker and placed in a wet chamber used for storing slides during all subsequent incubation steps. Specimens were incubated for 20 min with 2.5 % normal serum before gently tapping the slides sideways on paper towels to remove the serum. No washing step was performed at this stage. Each specimen was incubated for 60 min with the prepared solution of 2.5 % normal serum and primary antibody/antibodies at room temperature. Slides were washed for 2x 3 min in PBS containing 0.05 % Tween® 20. Specimens were then incubated for 60 min with the prepared solution of 2.5 % normal serum and secondary antibody/antibodies under aluminum foil at room temperature. The slides were washed in staining vessels covered with aluminum foil for 2x 3 min in PBS containing 0.05 % Tween® 20. Specimens were incubated for 60 min with the prepared solution of 2.5 % normal serum and fluorochrome-conjugated antibody/antibodies under aluminum foil at room temperature. Slides were washed in staining vessels covered with aluminum foil for 2x 3 min in PBS containing 0.05 % Tween® 20. The Vector TrueView Autofluorescence Quenching Kit (Biozol, VEC-SP-8400-15) was utilized according to the manufacturer’s instructions before washing slides in staining vessels covered with aluminum foil for 5 min in PBS containing 0.05 % Tween® 20. Finally, slides were mounted using VECTASHIELD Vibrance™ with DAPI Antifade Mounting Medium (Vector Laboratories/Biozol, VEC-H-1800), and coverslips were placed on top. Stained slides were digitized and analyzed using a ZEISS Axioscan 7 and the ZEN 3.5 (blue edition) software. The slides were stored in the dark at 4 °C.

#### Primary and Secondary Antibodies

All primary antibodies were pre-validated through repeated testing on appropriate tissue types using both immunohistochemistry (IHC) and immunofluorescence (IF) to ensure specific and robust staining. The primary antibodies used for IF and EvG-IHC staining of PPFE and CTRL samples are listed in Supplemental **Table E4**. Secondary antibodies for IF were similarly pre-evaluated in combination with well-characterized primary antibodies to confirm consistent performance

#### RNA *in-situ* hybridization (ISH)

RNA-ISH was conducted in adherence to the "RNAscope™ Multiplex Fluorescent Reagent Kit v2 User Manual" (UM 323100/Rev B/Effective Date: 10/11/2022) from ACD bio on FFPE lung explant tissue. Two to Three target probes were multiplexed during the assay in combination with conventional DAPI staining provided by the kit. The probes utilized are detailed in Supplemental Table E5. Post-staining, slides were preserved at 4°C in a dark environment until scanning. Digitalization and analysis of the stained slides were carried out using a ZEISS Axioscan 7 combined with ZEN 3.5 (blue edition) software.

#### Orcein and Elastica van Gieson stain

Formalin-fixed, paraffin-embedded (FFPE) tissue sections underwent standard deparaffinization and rehydration (1:30 min in xylene, followed by 1:30 min each in 100%, 90%, and 70% ethanol). Subsequently, German sections were incubated for 10 minutes in Resorcinol-Fuchsine solution according to Weigert, followed by differentiation in 100% ethanol (2 × 0:30 min). Iron hematoxylin staining (Weigert's method) was performed for 10 minutes, and sections were blued in tap water for 5 minutes. Slides were then counterstained in Van Gieson's solution (picrofuchsin) for 15 seconds. Meanwhile, French samples were stained with Orcein according to manufacturer instructions. Dehydration was carried out sequentially (0:05 min in 70% ethanol, 1:30 min in 90% ethanol, 2 × 1:30 min in 100% ethanol, and 1:30 min in xylene). Stained sections were digitized using a ZEISS Axioscan 7 and analyzed with ZEN 3.5 (blue edition) software. Slides were stored at room temperature.

#### Hierarchical phase-contrast tomography (HiP CT) Imaging

HiP CT imaging was performed on formalin inflated lung lobe (n=1[15]) as described before[16]. Virtual tissue cores were scanned within the intact inflated, formalin-fixed lung tissue. Raw images were processed with the neuroglancer pipeline (https://neuroglancer-docs.web.app/index.html). Pulmonary vein segmentation was performed on the zoomed HiP-CT data using an inhouse adopted nnU-Net framework. This framework has been trained on HiP-CT and micro-CT images at different resolutions,enabling it to handle diverse input data types. The resulting segmentation from the framework has been manually revised and corrected by a lung radiologist with +15 years of experience. To enhance visual clarity, the 3D visualization shows the main vessel structures from the highlighted region and was created in 3D Slicer 3.8.1.

#### Micro-CT Imaging

PPFE specimens were contrast-enhanced using either tungsten phosphoric acid (48 hours) or osmium tetraoxide contrastion (12h). In brief, the samples were first washed six times for 15 minutes each in cacodylate buffer. Post-fixation was carried out by incubating the samples in 1% osmium tetroxide for four hours on a rotator. Following fixation, the samples were rinsed twice for 15 minutes in cacodylate buffer. This was followed by eight washes of 10 minutes each in distilled water. The samples were then incubated overnight in 1% uranyl acetate. On the next day, they were washed twelve times for 10 minutes in cacodylate buffer. Dehydration was performed in a graded acetone series, consisting of four steps of 30 minutes each in 70% acetone, followed by four steps of 30 minutes each in 90% acetone, and finally six steps of 30 minutes each in 100% dry acetone. Contrasted formalin-fixed PPFE tissue specimen or native FFPE blocks were subsequently scanned with a Phoenix Nanotom® M micro-CT system (Waygate Technologies). Imaging was performed at a voxel resolution of 8.46 µm using an X-ray tube voltage of 60 kV and a current of 110 µA. Micrographs were captured with an average dynamic range of >2000 grey levels to ensure optimal contrast. Raw data underwent preprocessing, including region-of-interest (ROI) selection and inline median filtering, prior to 3D volume reconstruction using VGSTUDIO 2022 (64-bit, Volume Graphics).

#### Correlative Ultrastructure Analysis (CUA)

CUA analysis was carried out as recently described[17]. In brief, osmium tetraoxide contrasted OCT embedded tissue specimen was sectioned on a kryotstat (Leica, Wetzlar, German) in 50µm thick sections. Sections were subsequently fixed overnight with 1.5% paraformaldehyde, 1.5% glutaraldehyde in 0.15M HEPES buffer (pH 7.35). Afterwards, sections were postfixed in 1 % uranyl acetate (Serva, Heidelberg, Germany) over night and in 1% osmium tetroxide (EMS, Hatfield, PA) for 2 hours and after washing and dehydration in acetone embedded in Epon (Serva, Heidelberg, Germany). Ultrathin sections were imaged using a transmission electron microscope (model 364 Morgagni 268, FEI, Eindhoven, Netherlands). Final correlation of TEM images and light microscopic images was performed manually. Single obtained electron microscopic images were manually stitched together for improved parenchymal context.

#### Image Processing

Qualitative image analysis and processing of histological sections was carried out using ZEN (blue edition) software (ZEISS). For all immunofluorescence and RNA-ISH images, the "*background subtraction*" function was applied using default parameters. White and black point values were manually adjusted for each channel. No additional image processing techniques were employed. Multi-panel figure assembly was performed using Adobe Illustrator 2024.

#### Cell Segmentation and Mashine Learning guided quantification

For quantitative image analysis IF or ISH scans were loaded into QuPath (0.5.1)[18]. Cell segmentation was carried out based on DAPI stains employing the StarDist extension[19] following the recommended QuPath workflow referred to in the Docs ("https://qupath.readthedocs.io/en/stable/"). For StarDist based cell segmentation percentile normalization was applied using the 1st and 99th percentiles. A probability threshold of 0.5 was set for object detection. The analysis was conducted at a resolution of 0.5 µm per pixel. Detected nuclei were expanded by 5 µm to approximate whole-cell boundaries, with cell expansion constrained to 40% of the nuclear size to avoid overestimation. Shape descriptors and intensity measurements were calculated across all cellular compartments. Prediction probabilities were included as an additional measurement parameter. Segmented cell masks were subsequently employed for manual defining channel-wise training data for the QuPath inbuilt KNN classifier. App. 30-50 cells per channel per slide were manually annotated and KNN training was performed. Channel-related classifications were carefully reviewed and exported to .txt files. Zones of interest (i.e. Subpleural Fibrosis Zone, Elastofibrosis Zone, usual fibrotic niche) were manually segmented and annotated on the same slide in in agreement with two board certified pathologists with extensive experience in the histopathological diagnostics in ILDs. Area size was exported to R and cell densities per surface unit (µm$^2$) were calculated. Remodeled and un-remodeled parenchymal surface fractions of PPFE samples were likewise manually annotated on serial sectioned EvG or Orcein stained slides matched to the used 50µm slides employed for nuclei isolation.

#### Web Tool

The interactive dataset explorer was assembled with the R package ShinyCell[20].

### Supplementary Figure Legends

**Figure S1. HRCT Array.**

Overview of HRCT scans of enrolled PPFE patients in the German (A) and French cohort (B). HRCT are presented where available in coronar, sagittal and transversal projection. Anterior-posterior thorax diameters were assessed as demonstrated as the largest distance between sternal back and thorakal spine. Depth of suprasternal notching was assessed based on image qualities of available reformations either in coronar or sagittal projection.

**Figure S2. Array of selected assessed micro-CTs.**

Micro-CT assessment was conducted of supleural PPFE specimen either following contrasting with osmium-tetraoxide (A), tungsten phosphoric acid (B) or paraffin embedding (C). The dashed lines indicate the remodeling edge between AFE and alveolar parenchyma within the rendered 3D reconstructions. Vasculature was observed to rejuvenate from the alveolar to the elastofibrotic remodeled parenchyma (white arrows). Tungsten phosphoric acid contrast additionally revealed numerous thin vessels originating from the pleura (black arrows).

**Figure S3. Array of EvG and Orcein stained FFPE specimen.**

Overview of Elastica van Gieson (A) or Orcein (B) stained PPFE explant specimen of the German and French PPFE cohort. Serial sectioned 50µm thin sections were used for nuclei isolation for subsequent snRNAseq analysis.

**Figure S4. Gene overrepresentation analysis of DE genes of adventitial and elastofibrotic Fibroblasts.**

Clusters of enriched Reactome (v2024) terms based on MAST[13] related differentially expressed genes were generated by greedy clustering. Nodes denote counts of enriched genes, while edges represent the overlap among enrichment terms. Graphical representation was created with single cell pipeline package v0.5.6 (https://github.com/zhanghao-njmu/SCP).

**Figure S5. Multiplex stains of AEC1 and AEC2 located at the fibro-alveolar remodeling edge.**

Example stains of injured AEC1 marked by co-expression of AGER and CTSE (A) and intermediate AEC2 indicated by expression of SFTPC, CTSE and KRT17 (B).

**Figure S6. Average Expression of *CXCL12*, *CXCL14* and *PDGFRA* per Subject in Selected Celltypes.**

Gene expression is depicted as average per subject within each cohort. Statistical comparisons between PPFE-GER, PPFE-FR and CTRL were performed using Wilcoxon Rank-Sum test with *: $p \leq 0.05$, **: $p \leq 0.01$, ***: $p \leq 0.001$, and ****: $p \leq 0.0001$.

**Figure S7. Correlation of Annotated Parenchyma Fractions and Lineage Frequencies.**

Pearson correlation was performed between relative fractions of remodeled parenchyma and lineage frequencies. Correlations with pval<0.05 were considered statistically significant.

**Figure S8. Expression signatures of TLS related lymphocytes.**

Nebulosa plots of cells co-expressing indicated marker combinations. Each expression profile is depicted by a distinct color scale.

### Short Supplementary Table Legends

**Table S1. Summary of used primary antibodies.**

Clone and manufacturer information for the used primary antibodies.

**Table S2. Summary of used probes for RNA *in-situ* hybridization.**

A list of all used probes for validation.

**Table S3. Summary of used probes for RNA *in-situ* hybridization.**

Detailed clinical cohort statistics of the enrolled patient from both cohorts.

**Table S4. Overview of lineage related cell frequencies.**

descriptive statistics of average lineage frequencies per subject expressed as in mean±SD per cohort

**Table S5. Inter-Cohort Comparison of lineage frequencies**

Results of Wilcoxon-Rank sum test of lineage frequencies among cohorts corrected for multiple comparisons among lineages and cohorts with Bonferroni Correction.

**Table S6. Marker table of epithelial cell types.**

Results of Bonferroni-corrected Wilcoxon rank-sum test of each epithelial cell-type against the other

epithelial varieties, using the average gene expression per subject, per cell type including for multiple testing.

**Table S7. Overview of cell frequencies in the epithelial lineage.**

Descriptive statistics of average epithelial frequencies per subject expressed as in mean±SD per cohort

**Table S8. Inter-Cohort Comparison of Epithelial Celltypes**

Results of Wilcoxon-Rank sum test of epithelial cell frequencies among cohorts corrected for multiple comparisons among celltypes and cohorts with Bonferroni correction.

**Table S9. ssGSEA results of enrichment for GOBP 2021 related pathways**

Overview of ssGSEA related enrichment results for epithelial subcelltypes. Analysis was carried out under use of the escape R package (v 2.2.3).

**Table S10. Overview of cell frequencies in the mesenchymal lineage.**
Descriptive statistics of average mesenchymal cell frequencies per subject expressed as in mean±SD per cohort.

**Table S11. Inter-Cohort Comparison of mesenchymal celltype frequencies.**
Results of Wilcoxon-Rank sum test of mesenchymal cell frequencies among cohorts corrected for multiple comparisons among celltypes and cohorts with Bonferroni correction.

**Table S12. Marker table of mesenchymal celltypes.**
Results of Bonferroni-corrected Wilcoxon rank-sum test of each epithelial cell-type against the other epithelial varieties, using the average gene expression per subject, per cell type including for multiple testing.

**Table S13. Differential expressed genes among adventitial and elastofibrotic fibroblasts**
Overview of differentially expressed marker genes of adventitial vs. adventital-like fibroblasts. Analysis was performed on cell level.

**Table S14. Overview of cell frequencies in the endothelial lineage.**
Descriptive statistics of average endothelial cell frequencies per subject expressed as in mean±SD per cohort.

**Table S15. Inter-Cohort Comparison of endothelial celltype frequencies.**
Results of Wilcoxon-Rank sum test of endothelial cell frequencies among cohorts corrected for multiple comparisons among celltypes and cohorts with Bonferroni correction.

**Table S16. Marker table of mesenchymal celltypes.**
Results of Bonferroni-corrected Wilcoxon rank-sum test of each endothelial cell-type against the other endothelial varieties, using the average gene expression per subject, per cell type including for multiple testing.

**Table S17. Overview of lymphoid cell frequencies.**

Descriptive statistics of average lymphoid cell frequencies within all sampled nuclei per subject expressed as in mean±SD per cohort.

**Table S18. Inter-Cohort Comparison of lymphoid celltype frequencies.**

Results of Wilcoxon-Rank sum test of lymphoid cell frequencies among cohorts corrected for multiple comparisons among celltypes and cohorts with Bonferroni correction.

**Table S19. Overview of cell frequencies in the lymphoid lineage**

Descriptive statistics of average lymphoid cell frequencies per subject expressed as in mean±SD per cohort.

**Table S20. Marker table of lymphoid celltypes.**

Results of Bonferroni-corrected Wilcoxon rank-sum test of each lymphoid cell-type against the other lymphoid varieties, using the average gene expression per subject, per cell type including for multiple testing.

**Table S21. Overview of cell frequencies in the myeloid lineage.**

Descriptive statistics of average myeloid cell frequencies per subject expressed as in mean±SD per cohort.

**Table S22. Inter-Cohort Comparison of myeloid celltype frequencies.**

Results of Wilcoxon-Rank sum test of myeloid cell frequencies among cohorts corrected for multiple comparisons among celltypes and cohorts with Bonferroni correction.

**Table S23. Marker table of myeloid celltypes.**

Results of Bonferroni-corrected Wilcoxon rank-sum test of each myeloid cell-type against the other myeloid varieties, using the average gene expression per subject, per cell type including for multiple testing.

**Table S24. Top 2000 Ligand-Receptor Pairs by MultiNicheNetR**

Hierarchized overview of top 2000 ligand-receptor pairs within ranked by pseudobulk product of ligand in sender and receptor in receiver cell type.

**Table S25. Target genes of of endothelial JAM2, PDGFC and BMP2.**

Downstream Signaling in adventitial and elastofibrotic fibroblasts was predicted by signaling of JAM2, PDGFC and BMP2. Enrichment of target genes was performed in the GOBP database (v2021).

**Table S26. Enrichment analysis of significant fibroblast subtype related DE genes.**

Downstream enrichment DE genes was performed for Reactome terms (v3.16.0) using ClusterProfiler.

**Table S27. Greedy Clustering of enriched Reactome terms in Adventitial Fibroblasts.**

Clustering of enriched Reactome terms was performed using the greedy clustering.

**Table S28. Greedy Clustering of enriched Reactome terms in elastofibrotic Fibroblasts.**

Clustering of enriched Reactome terms was performed using the greedy clustering.

**Table S29. Pseudobulk Analysis (PPFE vs. CTRL) in the mesenchymal Lineage.**

Pseudobulk analysis was conducted with DESeq2 on subject level.

**Table S30. Pseudobulk Analysis (PPFE vs. CTRL) in alveolar fibroblasts.**

Pseudobulk analysis was conducted with DESeq2 on subject level.

**Table S31. DE Gene Enrichment Analysis of Pearson correlated Target Genes in signal receiving CTHRC1 fibrotic fibroblasts.**

Downstream Signaling in adventitial and *CTHRC1*+ fibrotic fibroblasts was predicted for Reactome pathways (v2024), using EnrichR.
