## Supplementary material for "The pleuroparenchymal fibroelastosis atlas reveals aberrant cell states and their zonation as an alternate roadmap to lung fibrosis": PPFE Supplements: PPFE_Supp_Figures_lowres_merged.pdf

A

*PPFE - GER*

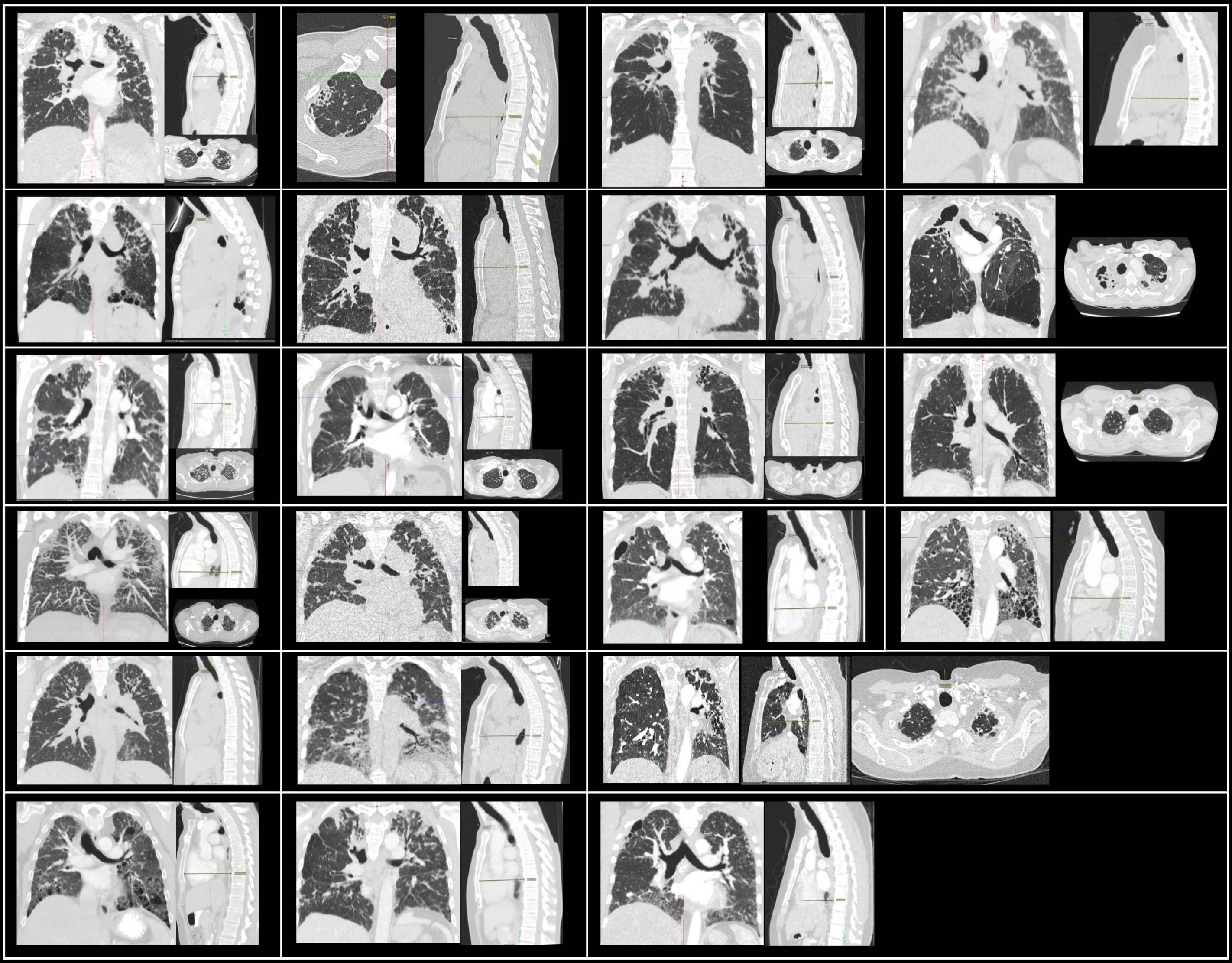

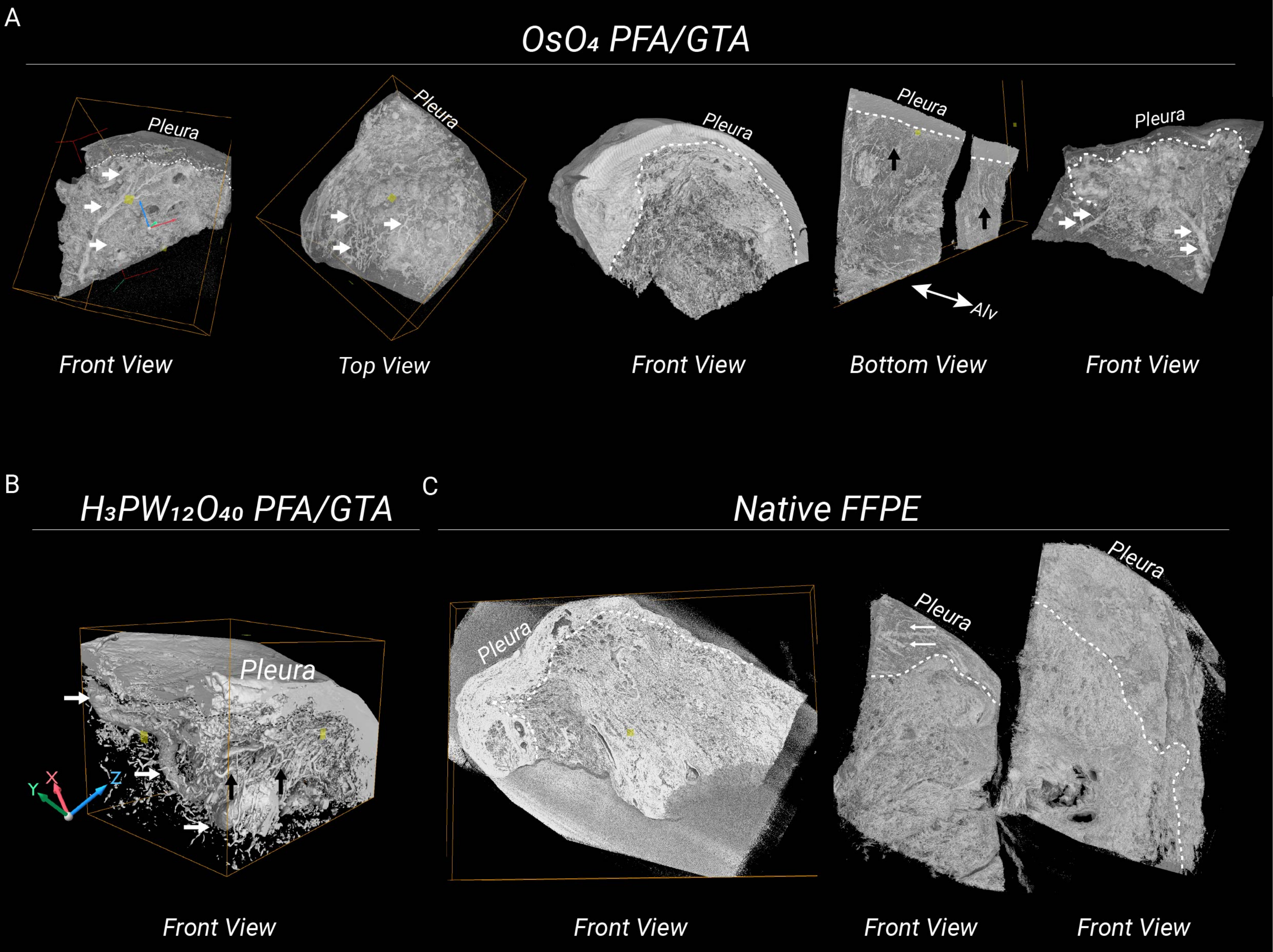
A
OsO₄ PFA/GTA
Pleura
Front View
Top View
Bottom View
Alv
B
H₃PW₁₂O₄₀ PFA/GTA
X
Y
Z
C
Native FFPE

A

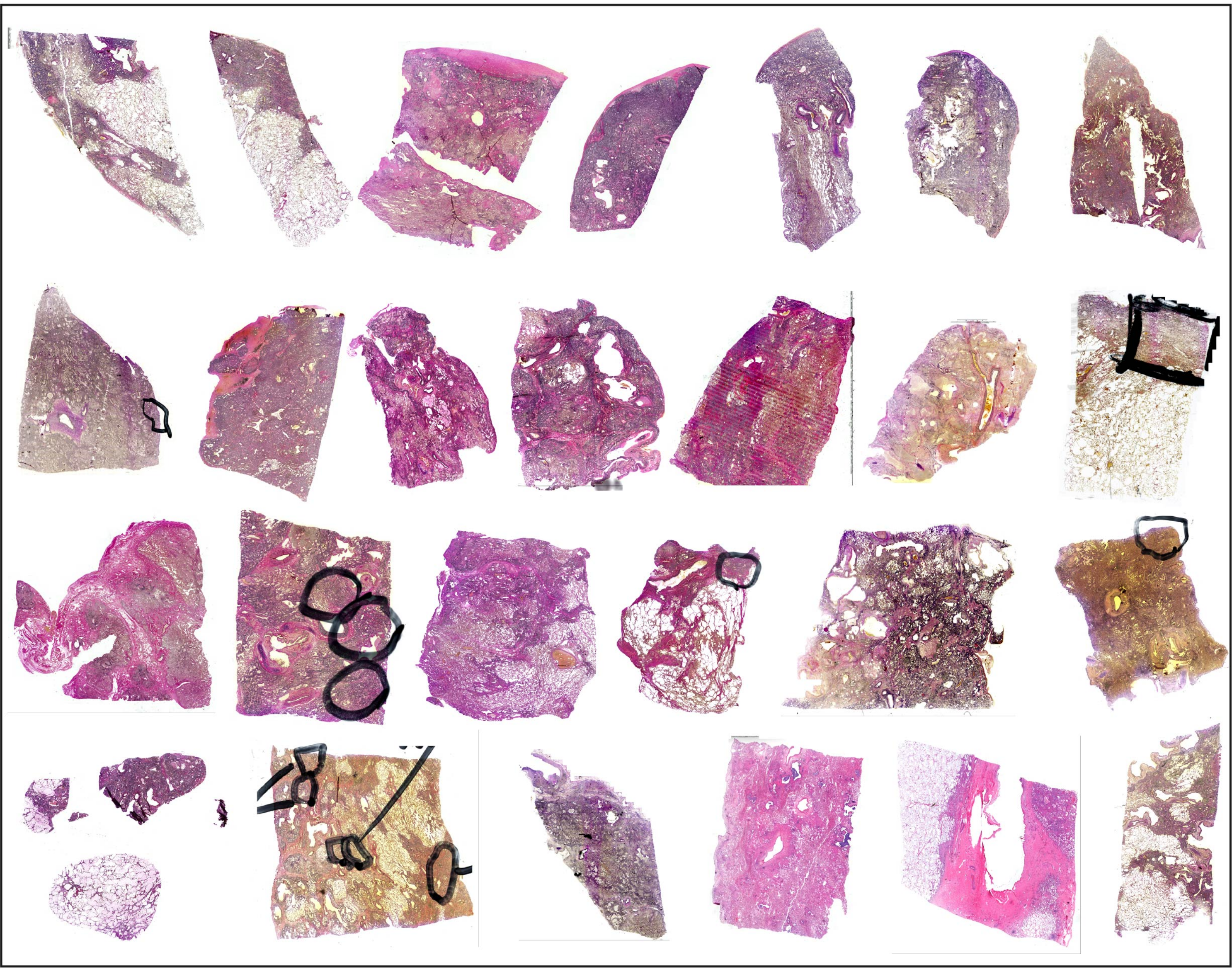
PPFE - GER (Elastica van Gieson Staining)

B

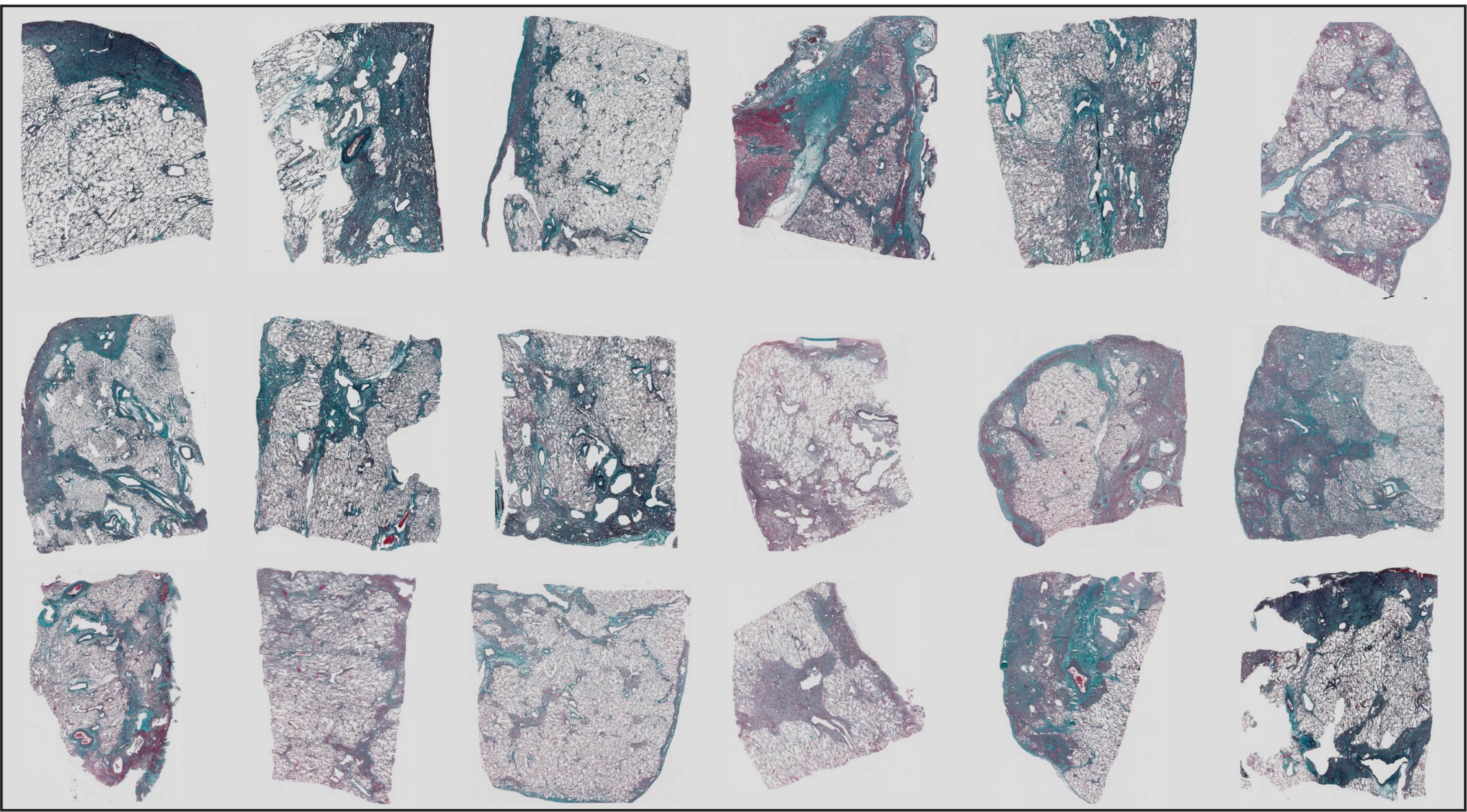
PPFE - FR (Orcein Staining)

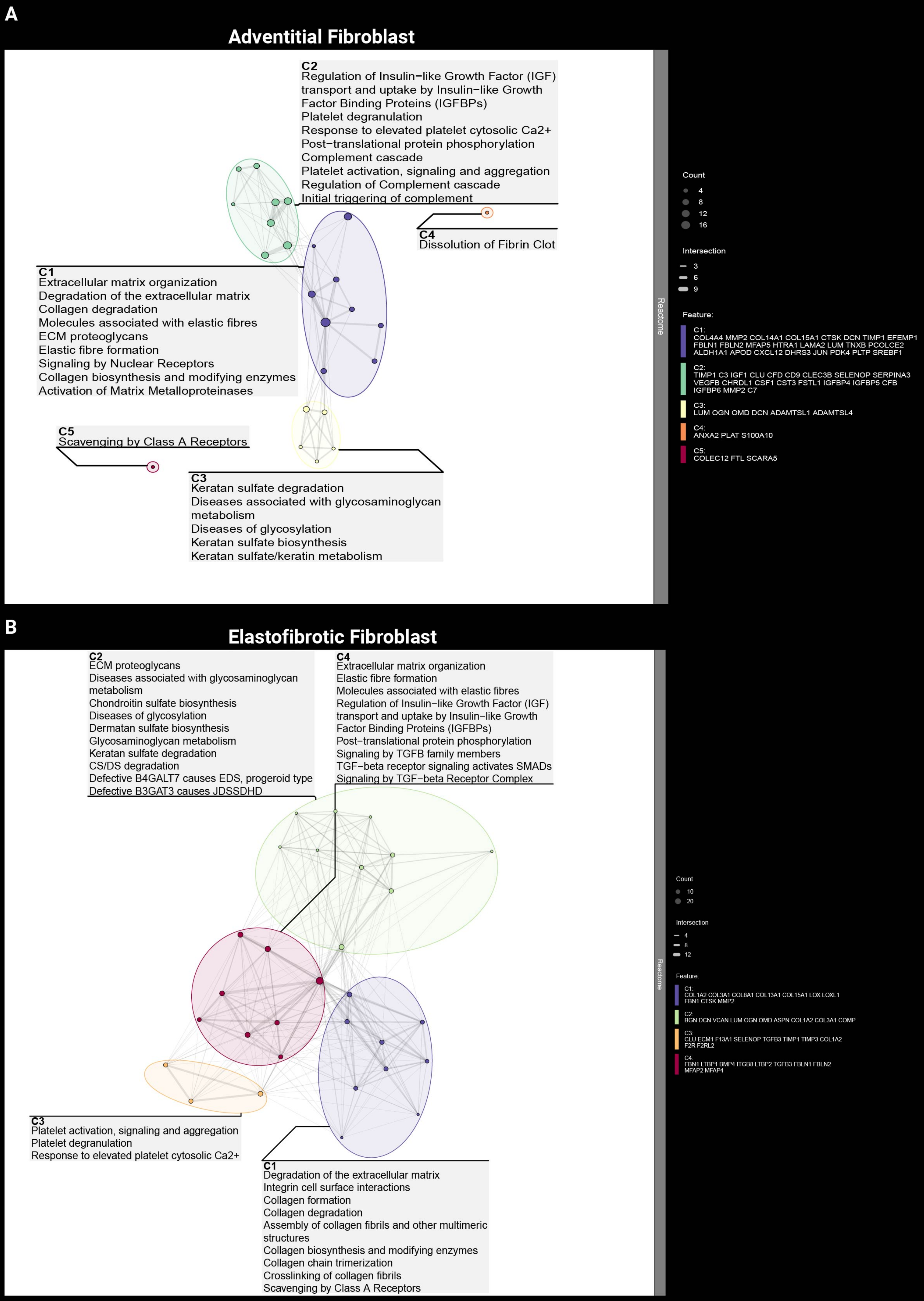
A
Adventitial Fibroblast
C2
Regulation of Insulin−like Growth Factor (IGF) transport and uptake by Insulin−like Growth Factor Binding Proteins (IGFBPs)
Platelet degranulation
Response to elevated platelet cytosolic Ca2+
Post−translational protein phosphorylation
Complement cascade
Platelet activation, signaling and aggregation
Regulation of Complement cascade
Initial triggering of complement
C4
Dissolution of Fibrin Clot
C1
Extracellular matrix organization
Degradation of the extracellular matrix
Collagen degradation
Molecules associated with elastic fibres
ECM proteoglycans
Elastic fibre formation
Signaling by Nuclear Receptors
Collagen biosynthesis and modifying enzymes
Activation of Matrix Metalloproteinases
C5
Scavenging by Class A Receptors
C3
Keratan sulfate degradation
Diseases associated with glycosaminoglycan metabolism
Diseases of glycosylation
Keratan sulfate biosynthesis
Keratan sulfate/keratin metabolism
Reactome
Count
4
8
12
16
Intersection
3
6
9
Feature:
C1:
COL4A4 MMP2 COL14A1 COL15A1 CTSK DCN TIMP1 EFEMP1 FBLN1 FBLN2 MFAP5 HTRA1 LAMA2 LUM TNXB PCOLCE2 ALDH1A1 APOD CXCL12 DHRS3 JUN PDK4 PLTP SREBF1
C2:
TIMP1 C3 IGF1 CLU CFD CD9 CLEC3B SELENOP SERPINA3 VEGFB CHRDL1 CSF1 CST3 FSTL1 IGFBP4 IGFBP5 CFB IGFBP6 MMP2 C7
C3:
LUM OGN OMD DCN ADAMTSL1 ADAMTSL4
C4:
ANXA2 PLAT S100A10
C5:
COLEC12 FTL SCARA5
B
Elastofibrotic Fibroblast
C2
ECM proteoglycans
Diseases associated with glycosaminoglycan metabolism
Chondroitin sulfate biosynthesis
Diseases of glycosylation
Dermatan sulfate biosynthesis
Glycosaminoglycan metabolism
Keratan sulfate degradation
CS/DS degradation
Defective B4GALT7 causes EDS, progeroid type
Defective B3GAT3 causes JDSSDHD
C4
Extracellular matrix organization
Elastic fibre formation
Molecules associated with elastic fibres
Regulation of Insulin−like Growth Factor (IGF) transport and uptake by Insulin−like Growth Factor Binding Proteins (IGFBPs)
Post−translational protein phosphorylation
Signaling by TGFB family members
TGF−beta receptor signaling activates SMADs
Signaling by TGF−beta Receptor Complex
C3
Platelet activation, signaling and aggregation
Platelet degranulation
Response to elevated platelet cytosolic Ca2+
C1
Degradation of the extracellular matrix
Integrin cell surface interactions
Collagen formation
Collagen degradation
Assembly of collagen fibrils and other multimeric structures
Collagen biosynthesis and modifying enzymes
Collagen chain trimerization
Crosslinking of collagen fibrils
Scavenging by Class A Receptors
Reactome
Count
10
20
Intersection
4
8
12
Feature:
C1:
COL1A2 COL3A1 COL8A1 COL13A1 COL15A1 LOX LOXL1 FBN1 CTSK MMP2
C2:
BGN DCN VCAN LUM OGN OMD ASPN COL1A2 COL3A1 COMP
C3:
CLU ECM1 F13A1 SELENOP TGFB3 TIMP1 TIMP3 COL1A2 F2R F2RL2
C4:
FBN1 LTBP1 BMP4 ITGB8 LTBP2 TGFB3 FBLN1 FBLN2 MFAP2 MFAP4

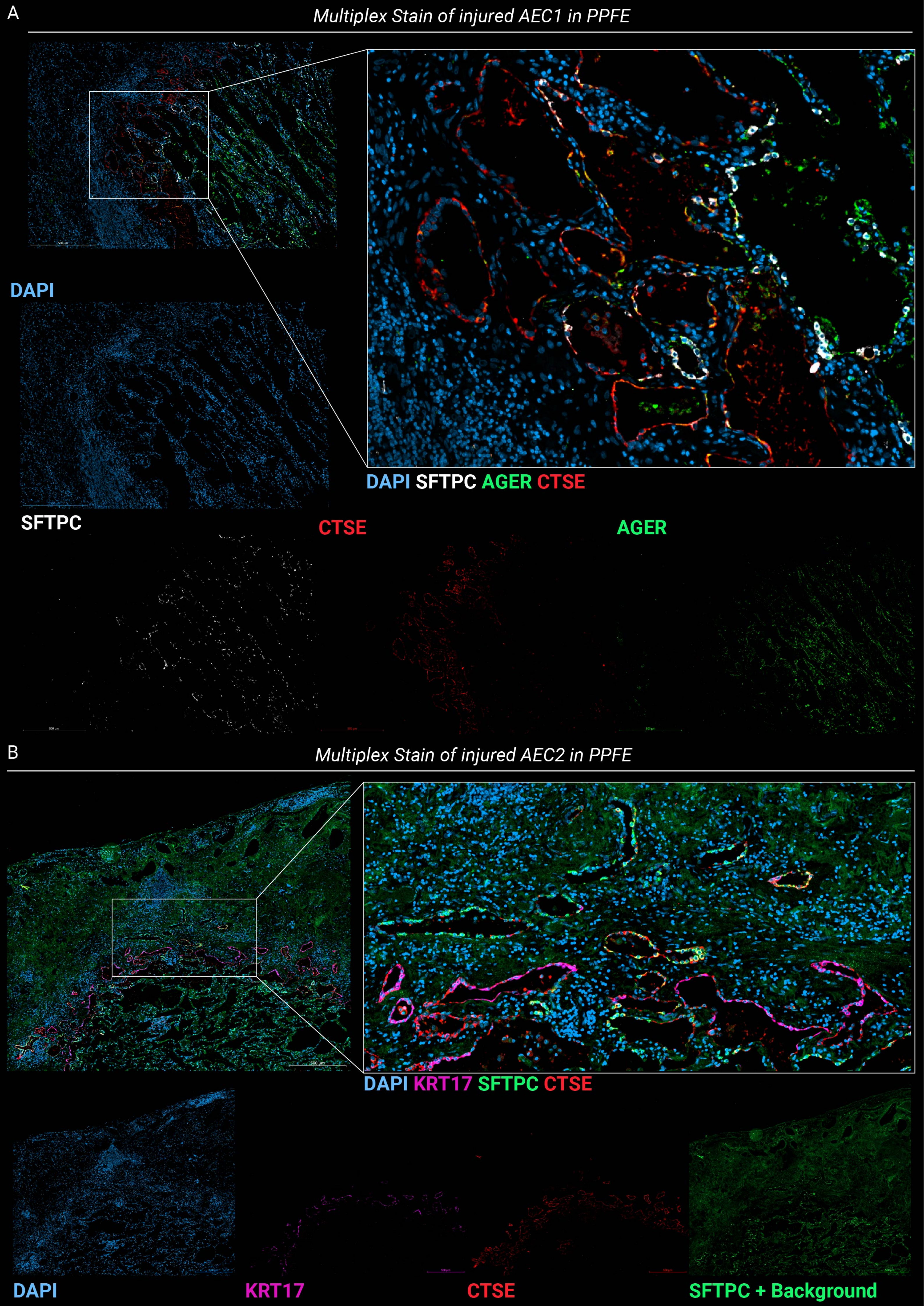
A
Multiplex Stain of injured AEC1 in PPFE
DAPI
DAPI SFTPC AGER CTSE
SFTPC
CTSE
AGER
B
Multiplex Stain of injured AEC2 in PPFE
500 µm
DAPI KRT17 SFTPC CTSE
DAPI
KRT17
CTSE
SFTPC + Background

### Average Expression per Subject in selected Cell Types

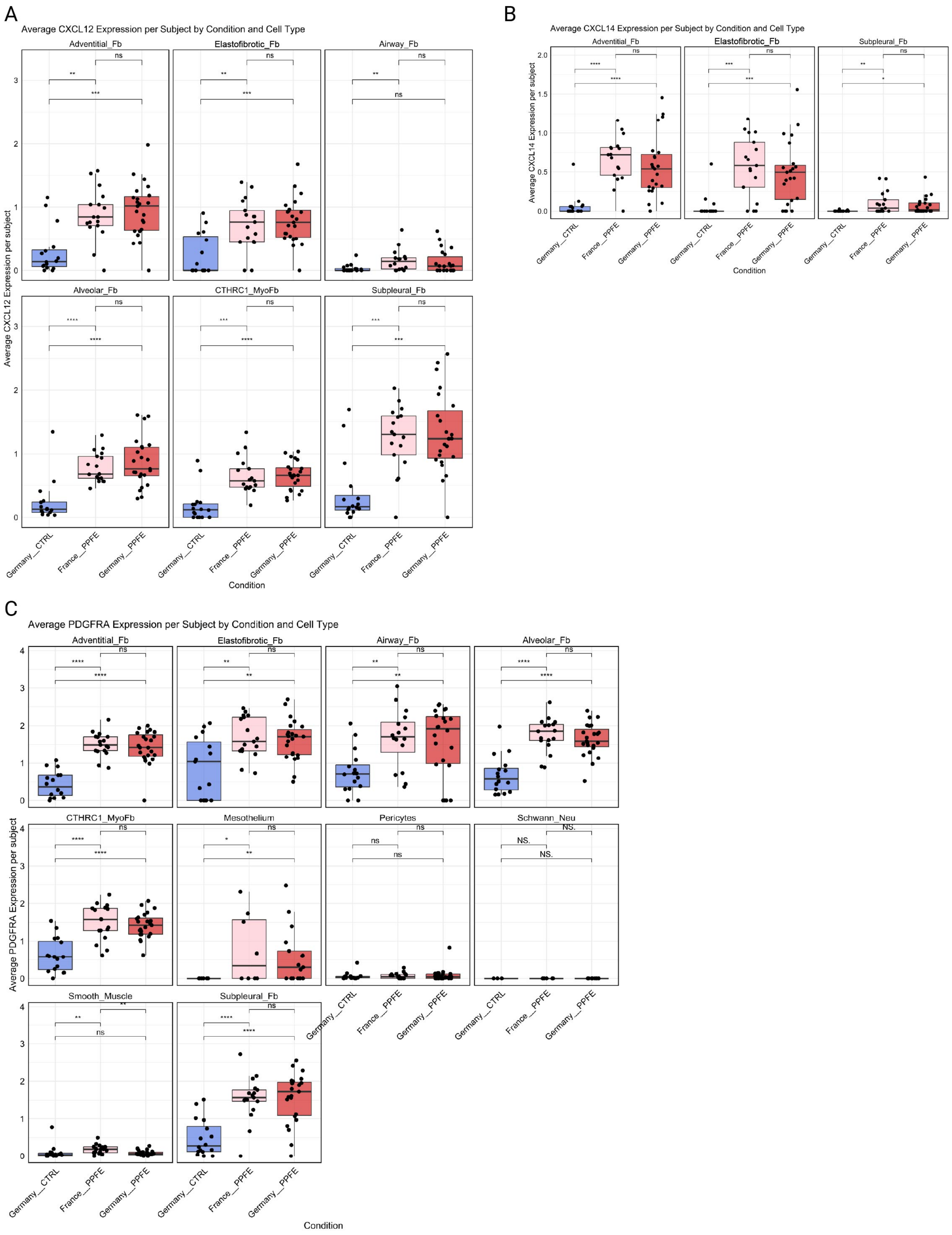

A

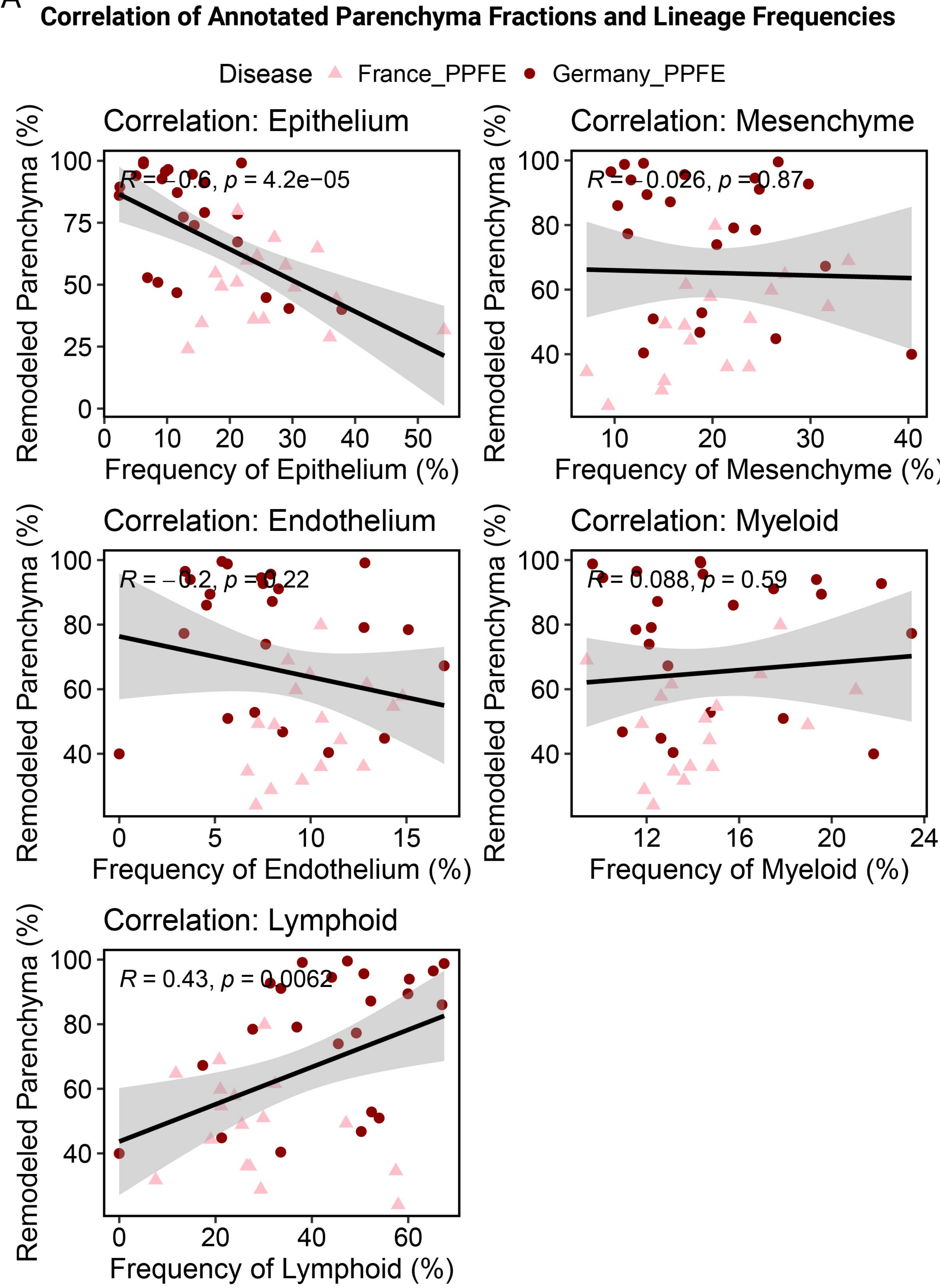
Correlation of Annotated Parenchyma Fractions and Lineage Frequencies
Disease France_PPFE Germany_PPFE
Correlation: Epithelium
R = −0.6, p = 4.2e−05
Remodeled Parenchyma (%)
Frequency of Epithelium (%)
Correlation: Mesenchyme
R = −0.026, p = 0.87
Remodeled Parenchyma (%)
Frequency of Mesenchyme (%)
Correlation: Endothelium
R = −0.2, p = 0.22
Remodeled Parenchyma (%)
Frequency of Endothelium (%)
Correlation: Myeloid
R = 0.088, p = 0.59
Remodeled Parenchyma (%)
Frequency of Myeloid (%)
Correlation: Lymphoid
R = 0.43, p = 0.0062
Remodeled Parenchyma (%)
Frequency of Lymphoid (%)

### TLS Related Lymphocyte Cell Types

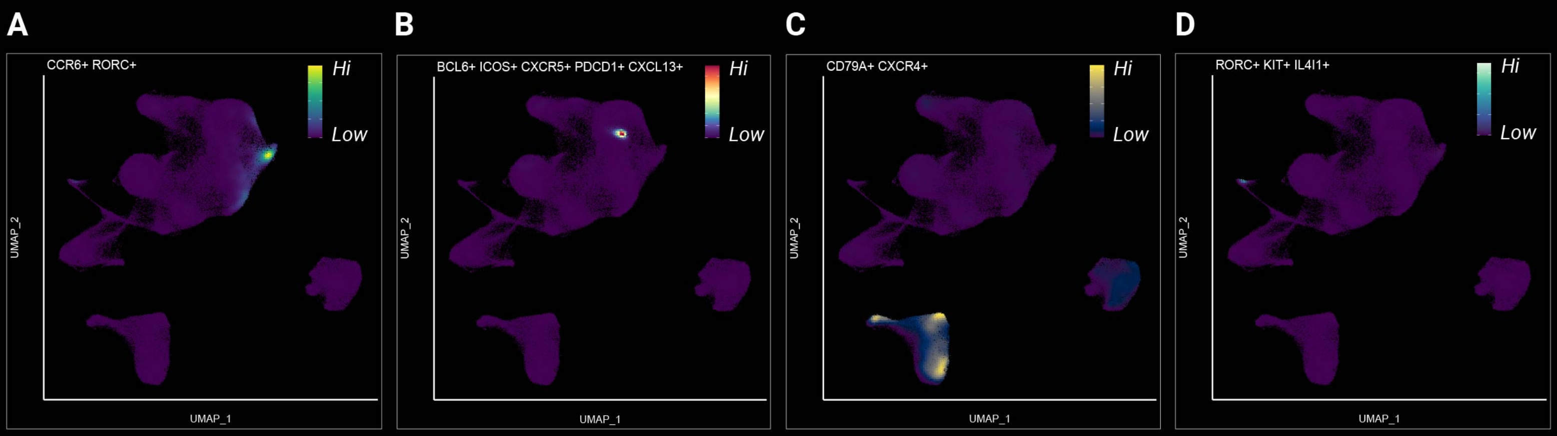
